# Complement C5a impairs phagosomal maturation in the neutrophil through phosphoproteomic remodelling

**DOI:** 10.1101/2020.01.17.907618

**Authors:** Alexander J.T. Wood, Arlette M. Vassallo, Marie-Hélène Ruchaud-Sparagano, Jonathan Scott, Carmelo Zinnato, Carmen Gonzalez-Tejedo, Kamal Kishore, Clive S. D’Santos, A. John Simpson, David K. Menon, Charlotte Summers, Edwin R. Chilvers, Klaus Okkenhaug, Andrew Conway Morris

## Abstract

Critical illness is accompanied by the release of large amounts of the anaphylotoxin, C5a. C5a suppresses antimicrobial functions of neutrophils which is associated with adverse outcomes. The signalling pathways that mediate C5a-induced neutrophil dysfunction are incompletely understood. Healthy donor neutrophils exposed to purified C5a demonstrated a prolonged defect (7 hours) in phagocytosis of *Staphylococcus aureus*. Phosphoproteomic profiling of 2712 phosphoproteins identified persistent C5a signalling and selective impairment of phagosomal protein phosphorylation on exposure to *S. aureus.* Notable proteins included early endosomal marker ZFYVE16 and V-ATPase proton channel component ATPV1G1. A novel assay of phagosomal acidification demonstrated C5a-induced impairment of phagosomal acidification which was recapitulated in neutrophils from critically ill patients. Examination of the C5a-impaired protein phosphorylation indicated a role for the phosphatidylinositol 3-kinase VPS34 in phagosomal maturation. Inhibition of VPS34 impaired neutrophil phagosomal acidification and killing of *S. aureus*. This study provides a phosphoproteomic assessment of human neutrophil signalling in response to *S. aureus* and its disruption by C5a, identifying a defect in phagosomal maturation and new mechanisms of immune failure in critical illness.

## Introduction

Critically ill patients who require exogenous organ support as a result of severe physiologic insult, are at high risk of secondary infections (Vincent *et al*, 2009). Critical illness may arise from a variety of sterile or infectious insults. However, despite its varied aetiology, critical illness is often accompanied by stereotyped immune dysregulation, with features of both hyperinflammation and immune-mediated organ damage, as well as impairment of anti-microbial functions (Meakins *et al*, 1977; Conway Morris *et al*, 2013; Hotchkiss *et al*, 2013a). Critical illness is estimated to cause 58 million adult deaths per year globally, (Adhikari *et al*, 2010) and whilst much of the mortality is attributable to the underlying condition, secondary infections make a significant contribution to the eventual outcome (Adhikari *et al*, 2010; Scicluna *et al*, 2015; van Vught *et al*, 2016; Vincent *et al*, 2006).

Impairment of immune cell function predicts secondary infection, (Hotchkiss *et al*, 2013b; Conway Morris *et al*, 2013; Demaret *et al*, 2015; Landelle *et al*, 2010) and failure of neutrophil phagocytosis and bacterial killing has been demonstrated to be one of the strongest predictors of these infections. A key driver of the functional impairment of neutrophils is the anaphylatoxin C5a (Conway Morris *et al*, 2009, 2011; Huber-Lang *et al*, 2002b). However, there remain no efficacious treatments for critical-illness induced immune dysfunction, in part because the mechanisms that underpin C5a-induced dysfunction are incompletely understood.

A wealth of data have demonstrated the importance of C5a in driving classical inflammatory events in neutrophils, including chemotaxis (Ward & Newman, 1969; Ehrengruber *et al*, 1994), generation of reactive oxygen species (ROS) (Suire *et al*, 2006; Mazaki *et al*, 2006; Huber-Lang *et al*, 2002b), phagocytosis (Mollnes *et al*, 2002; Brekke *et al*, 2007), degranulation (Denk *et al*, 2017a, 2017b), and delayed apoptosis (Lee *et al*, 2008; Perianayagam *et al*, 2002, 2004). One of the key questions in this area is how to explain the divergent findings of C5a being a critical co-factor in phagocytosis (Mollnes *et al*, 2002; Brekke *et al*, 2007), but also capable of suppression of phagocytosis in sepsis and critical illness (Conway Morris *et al*, 2009, 2011; Huber-Lang *et al*, 2002b). In critical illness, dysregulated activation of the complement and coagulation cascades occurs, leading to exposure of neutrophils to high concentrations of C5a (Hotchkiss *et al*, 2013a; Venet & Monneret, 2018; Lord *et al*, 2014; Conway-Morris *et al*, 2018; Ward, 2004). In these circumstances, we and others have shown that C5a reduces neutrophil phagocytosis and ROS production in both rodent models and critically ill patients (Conway Morris *et al*, 2009, 2011; Czermak *et al*, 1999; Huber-Lang *et al*, 2002b). Further, C5a exposure has been shown to be associated with nosocomial infection, organ failure, and increased mortality in critically ill patients (Conway Morris *et al*, 2011, 2009, 2013; Czermak *et al*, 1999; Huber-Lang *et al*, 2001, 2002a).

Whilst several signals mediating aspects of C5a-induced neutrophil dysfunction have been established (Conway Morris *et al*, 2011; Denk *et al*, 2017a; Huber-Lang *et al*, 2002b), a global picture of signalling in neutrophils encountering common pathogens and how this process is perturbed by C5a does not exist. Such studies are challenging in neutrophils owing to their high degradative enzyme content and short *in-vitro* survival times (Luerman *et al*, 2010).

This study aimed to characterise the neutrophil phosphoprotein response to a common nosocomial pathogen, *Staphylococcus aureus*, and investigate how this is perturbed by prior exposure to C5a. Our differential phosphoprotein analysis implicated C5a in altered phagosomal maturation, findings that we confirmed with functional neutrophil assays in C5a-treated healthy donor cells and those from critically ill patients. The phosphoprotein response to *S. aureus* implicated the involvement of the phosphatidylinositol 3-kinase VPS34, hence we continued examined the effects of this enzyme on phagosomal maturation.

## Results

### C5a induces a prolonged defect in neutrophil phagocytosis of bacteria

C5a induces a defect in phagocytosis of the clinically relevant bacterial species *S. aureus* (Figure 1A) and *E. coli* (1B). Pulse exposure of neutrophils to C5a revealed a persistent defect in phagocytosis lasting at least seven hours (1C), with short pulses inducing a significant defect. These effects were not explained by the loss of cell viability (1D). A similar prolonged defect was identified in the whole blood assay (1E), representing continuous exposure of neutrophils to C5a (which cannot be washed off in this assay). The ability of C5a to inhibit phagocytosis was dependent on the temporal relationship between C5a and *S. aureus* exposure. Only pre-exposure to C5a induced the defect in phagocytosis, whereas co-exposure or the addition of C5a 30 minutes after *S. aureus* addition failed to induce a defect (1F). In a dose titration we could not identify any dose which enhanced phagocytosis, with progressive decrease in phagocytosis as concentration of C5a was increased (Figure S1A).

**Figure 1:**
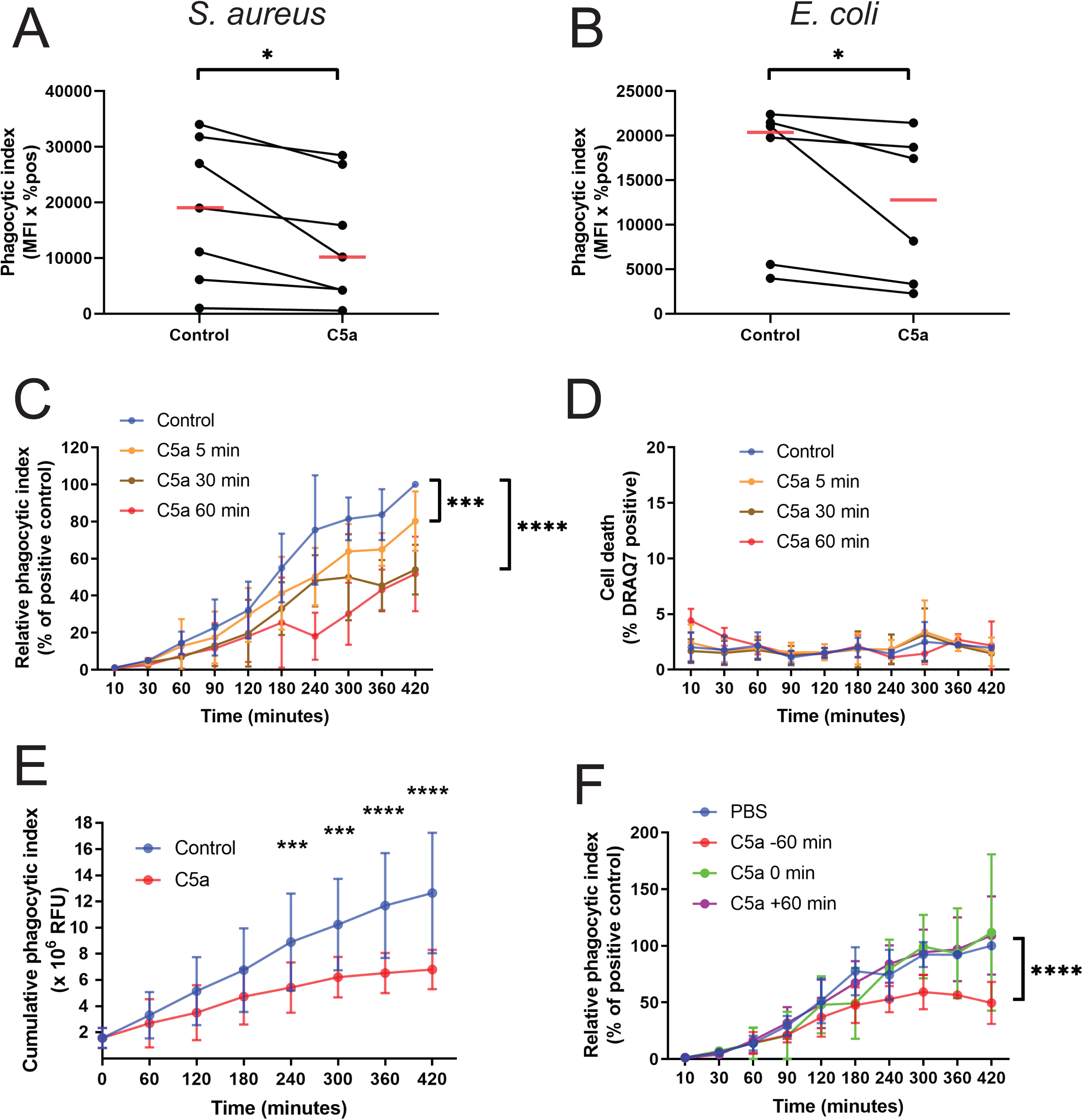
C5a induces a prolonged defect in neutrophil phagocytosis of bacteria. **A** and **B**: Isolated neutrophils were pre-treated with 100 nM C5a or vehicle control for 60 min before incubation with *S. aureus* (A) or *E. coli* (B) bioparticles. Data are presented as the median phagocytic index for each condition for n=7 (A) or 6 (B) independent donors, \**P* = 0.016 (A) and 0.031 (B) by Wilcoxon’s matched-pairs signed rank test. **C**: Neutrophils were pulsed with 100 nM C5a or PBS control for the indicated periods of time, followed by 2 washes. *S. aureus* bioparticles were then added and cells were incubated for the indicated time points. Data are presented as the mean and SD of the phagocytic index of C5a-treated cells relative to their paired vehicle control for n=5 independent experiments. *P* < 0.0001 for time and *P* = 0.0186 for treatment by two-way ANOVA. *\*\*\*P* = 0.0001 \*\*\*\**P* < 0.0001 by Dunnett’s multiple comparison test. **D**: Data are presented as the mean and SD of the percentage of DRAQ7 positive, dead cells for n=5 independent experiments. *P* = 0.378 for time *and P* = 0.349 for treatment by two-way ANOVA. **E**: Anticoagulated whole blood was pre-treated with 300 nM C5a or control for the indicated duration before phagocytosis was measured as previously indicated. Data are presented as the mean and SD of the cumulative phagocytic index for 4 independent experiments. *P* < 0.0001 by two-way ANOVA, \*\*\*\**P* < 0.0001, \*\*\**P* < 0.001 by Sidak’s multiple comparisons test. **F**: *S. aureus* particles were incubated with isolated PMNs in the presence of 100 nM C5a or PBS added at the indicated time points, with time 0 representing the time of addition of *S. aureus* bioparticles. Experiments proceeded for the indicated time points and phagocytic index quantified. Data are presented as the mean and SD of the phagocytic index of C5a-treated cells relative to their paired vehicle control for n=5 independent experiments. *P* < 0.0001 for time and *P* = 0.0186 for treatment by two-way ANOVA. \*\*\*\**P* < 0.0001 by Dunnett’s multiple comparisons test.

To explore the potential mechanisms whereby pre-exposure to *S. aureus* prevents the inhibitory effect of C5a, we examined whether this could be due to reduced C5aR1 expression. Although we could demonstrate a reduction in C5aR1 following *S. aureus* exposure (Figure S1B), this was modest and similar to the reductions induced by other inflammatory mediators including lipopolysaccharide (LPS) and leukotriene A (LTA), neither of which ameliorated the subsequent suppressive effect of C5a (Figure S1C). Further, C5a and not LPS, LTA, granulocyte-macrophage colony-stimulating factor (GM-CSF) and tumour necrosis factor (TNF) reduced neutrophil phagocytosis (Figure S1D). To confirm the functional relevance of C5a-impaired phagocytosis, we demonstrated that C5a pre-treatment reduced bacterial killing of *S. aureus* (Figure S1E).

Given the temporal dependence of the inhibitory effect of C5a, we examined whether prior exposure to *S. aureus* enhanced or impaired subsequent ingestion. Using sequential exposure of whole blood to *S. aureus* labelled with two different pHrodo probes, we demonstrated that neutrophils which ingest the first target are more likely to ingest the second target (Figure S2A), whilst progressive increase in the first target revealed a fundamental limit on the capacity of neutrophils to ingest (Figures S2B and C).

### S. aureus and C5a induce widespread changes in the neutrophil phosphoproteome

Although key signalling ‘nodes’ have been identified in neutrophils following C5a exposure (Conway Morris *et al*, 2009, 2011), no map of global signalling networks has been produced. Given the rapidity of the C5a-induced phagocytic impairment demonstrated above, and the known signalling kinetics of G-protein coupled receptors (GPCRs) (Lohse *et al*, 2008), we examined post-translational modification by phosphorylation (i.e. a phosphoproteomic approach).

In total, 4859 proteins and 2712 phosphoproteins were identified in peripheral blood neutrophils obtained from four healthy volunteers. C5a-induced suppression of phagocytosis in these donors was confirmed (Figure S3A), and technical reproducibility was high (Figures S3B-E) with the magnitude of phosphorylation changes within the previously reported range (Papachristou *et al*, 2018). Changes in the human proteome were minimal (2 % of total proteome with *S. aureus* treatment) whereas phosphoprotein expression varied markedly (31.6 % of total phosphoproteome with *S. aureus* treatment, Table S1). Figure 2 shows the top 2.5% most variable phosphoproteins with protein identification, whereas the top 25 % are shown in Figure S4 to demonstrate wider changes within the phosphoproteome. The phosphoproteomic and proteomic datasets are publicly available in the PRIDE database (data available to reviewers, will be made public on acceptance of manuscript).

**Figure 2:**
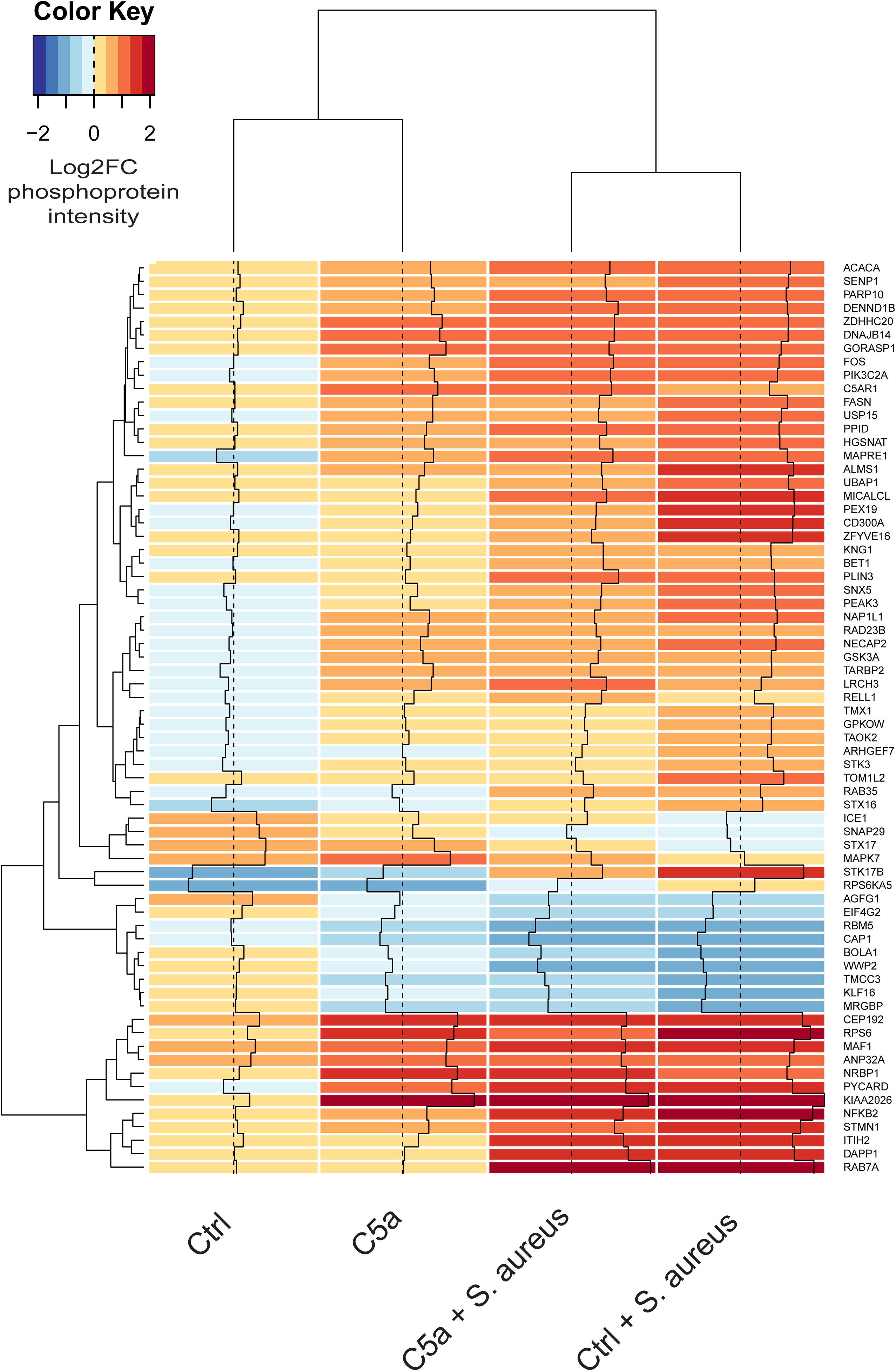
*S. aureus* and C5a induce widespread changes in the neutrophil phosphoproteome. Heatmap of phosphoprotein intensity relative to baseline (log_2_ fold change) across the four experimental conditions shows phosphoproteins with variance across conditions in the top 97.5^th^ centile with dendrograms clustered by Euclidean distance. Increased phosphoprotein expression is indicated in red, decreased in blue. Only phosphoproteins detected in all four donor samples were included.

### C5a exposure induces persistent alteration in phosphoproteins across several pathways

Figure 3A shows a volcano plot comparing neutrophils treated with C5a versus vehicle control. 119 proteins were significantly differentially phosphorylated at 1 hour, indicating persistent signalling, consistent with the prolonged inhibition of phagocytosis seen in Figure 1. Notably, C5aR1 remained highly phosphorylated (a modification key to its internalisation) (Braun *et al*, 2003) and this change has been used to identify C5a-exposed, dysfunctional neutrophils (Conway Morris *et al*, 2009, 2011, 2013, 2018; Schmidt *et al*, 2015; Unnewehr *et al*, 2013). Pathway enrichment using Metascape (Zhou *et al*, 2019) indicated involvement of pathways including membrane trafficking, regulated exocytosis (degranulation), and phosphatidylinositol-3,4,5-trisphosphate (PIP3) signalling which persist one hour after stimulation with C5a (Figure 3B). Given the finding of a temporal dependency of C5a exposure on phagocytosis (Figure 1F), we examined how the phosphoproteomic alterations after C5a compared to those after *S. aureus.* The pathways showing prolonged activation after C5a exposure overlap with those induced by *S. aureus* (Figure S5A) and many of the proteins are common to both conditions (Figure S5B).

**Figure 3:**
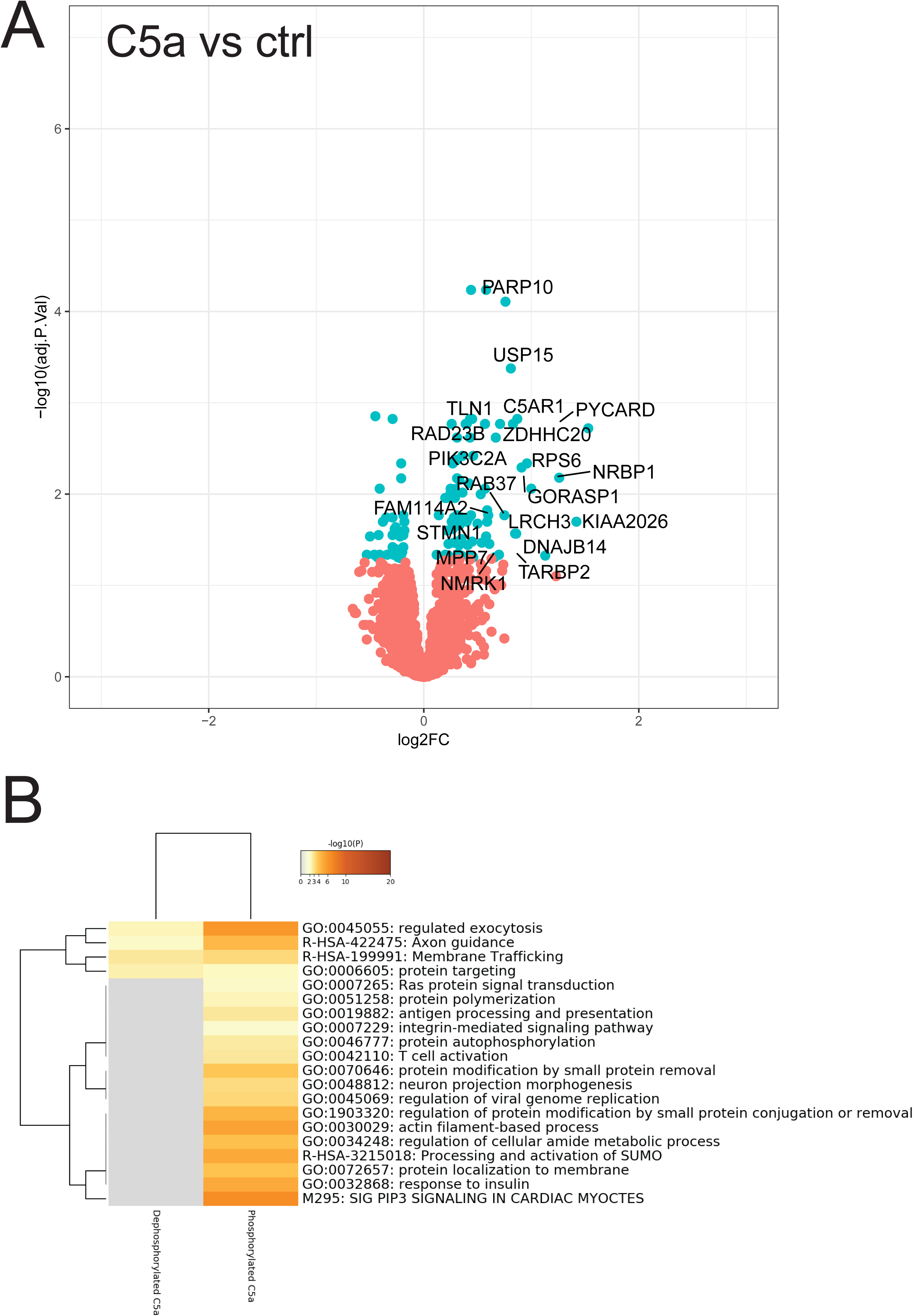
C5a exposure induces persistent alteration in phosphoproteins across several pathways. **A:** Proteins with adjusted *P*-values < 0.05 are shown in blue and the 20 proteins with the highest absolute log_2_ fold change are labelled. *P*-values were computed by limma-based linear models with Bonferroni’s correction for multiple testing. **B:** Metascape(Zhou *et al*, 2019) enrichment heatmap showing functional clusters of phosphoproteins affected by C5a treatment.

### *S. aureus* induces a marked alteration in the phosphoproteome which is significantly impacted by C5a exposure

Exposure of neutrophils to *S. aureus* induced a marked alteration in the phosphoproteome (Figure 4A); 863 proteins (31% of the phosphoproteome) significantly alter their phosphor-status. Pathway enrichment indicated the involvement of multiple pathways, notably Rho-GTPase signalling, endosomal transport, degranulation, and actin cytoskeleton organisation (Figure 4B, with extended heatmap showing top 100 pathways shown in Figure S6).

C5a exposure prior to *S. aureus* reduced the phosphoprotein response to the bacterium considerably (Figure 4C). However, comparing C5a and control treated cells exposed to *S. aureus*, 19 proteins were identified, suggesting selective pathway modulation (Figure 4D). When mapped to known pathways using Metascape (Zhou *et al*, 2019) and manually annotated from the Uniprot database (The Uniprot Consortium, 2019), a pattern of reduced phosphorylation of phagosomal maturation proteins (Table 1) and pathways (Figure 4E) emerged. Notably, early endosomal marker ZFYVE16 and its interactor TOM1 had impaired phosphorylation following C5a exposure, as did V-type ATPase subunit G1 (which is critical for phagosomal acidification). ZFYVE16 requires phosphatidylinositol-3-phosphate (PI3P) for recruitment to the phagosome (Sorkin & Von Zastrow, 2009). Another prominent PI3P-responsive protein noted was Ras-related protein 7a (RAB7A), although this protein was not differentially phosphorylated between the C5a/*S. aureus* and vehicle control/*S. aureus* conditions. Figure S7 shows individual donor data for these key proteins.

**Table I:** Differentially phosphorylated proteins between C5a and control-treated neutrophils exposed to *S. aureus*. All 19 phosphoproteins with Bonferroni adjusted p-values < 0.05 for difference in phosphorylation status between the Control plus *S. aureus* vs C5a plus *S. aureus* conditions. Subcellular location and function manually annotated from Uniprot database (The Uniprot Consortium, 2019).

| Protein name | Sub-cellular localisation | Functional role |
| --- | --- | --- |
| **Endosomal/phagosomal** |  |  |
| Zinc finger FYVE domain-containing protein 16 (ZFYVE16) | Early Endosomes | PI(3)P binding FYVE domain, found on early endosomes, targets TOM1 to endosomes |
| Target of Myb protein 1 (TOM1) | Endosomes/azurophil granule membrane (neutrophil) | Known interactor with ZFYVE16 |
| V-type proton ATPase subunit G 1 (ATP6V1G1) | Endosomes/phagosomes | Proton channel acidifying vacuolar structures |
| Wiskott-Aldrich protein (WAS) | Cytoplasm, phagosomes | RhoGTPase effector and actin polymerisation |
| Epsin-1 (EPN1) | Plasma membrane, clathrin coated pits | Regulates receptor-mediated endocytosis, PI(4,5)P binding |
| **Lysosomes** |  |  |
| Ubiquitin carboxyl-terminal hydrolase 5 (USP5) | Lysosome and cytoplasm | Deubiquitination |
| Synaptic vesicle membrane protein VAT-1 homolog (VAT1) | Lysosomes/azurophil granule (neutrophil) | Neutrophil degranulation |
| **Nuclear** |  |  |
| Nuclear pore complex protein 153 (NUP153) | Nuclear membrane | Phosphorylation occurs during nuclear membrane disassembly |
| Lamin-B1 (LMNB1) | Nuclear membrane | Phosphorylation occurs during nuclear membrane disassembly |
| Histone H3.3 (H3F3A) | Nucleus | Histone protein |
| Protein SGT1 homolog (SUGT1) | Nucleus and kinteochores | Mitosis |
| Sperm-associated antigen 7 (SPAG7) | Nucleus | Nucleic acid binding |
| Splicing factor U2AF 35 kDa subunit (U2AF1) | Nucleus | RNA splicing |
| Chromatin complexes subunit BAP18 (BAP18) | Nucleus | Chromatin organisation |
| **Nuclear/Cytoplasmic proteins** |  |  |
| Serine/threonine-protein kinase 3 (STK3) | Nucleus/cytoplasm | Cycles between nucleus and cytoplasm, influenced by phosphorylation status under control of caspases |
| Transforming acidic coiled-coil-containing protein 1 (TACC1) | Nucleus, centrosome, microtubules | Microtubule organisation and mitotic spindle formation |
| Serine/threonine-protein kinase TAO2 (TAOK2) | Nucleus/Cytoskeleton | Regulation of actin cytoskeleton and MAP kinase activity |
| **Endoplasmic reticulum** |  |  |
| Protein lunapark (LNP) | Endoplasmic reticulum | Phosphorylated during mitosis |
| Thioredoxin-related transmembrane protein 1 (TMX1) | Endoplasmic reticulum | Cell response to oxidative stress |

**Figure 4:**
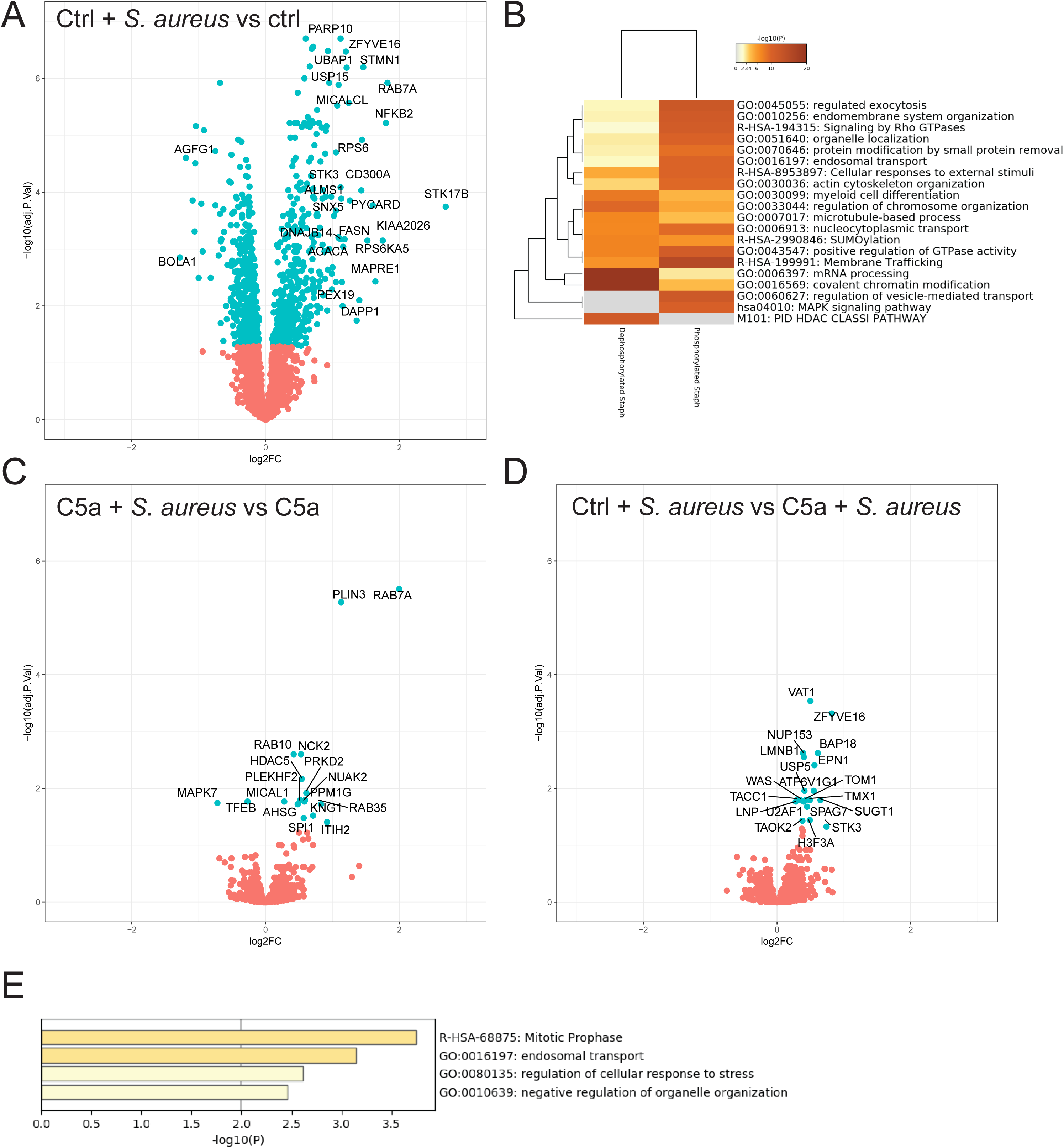
*S. aureus* induces a marked alteration in the phosphoproteome that is significantly impacted by C5a exposure. **A, B, D:** Proteins with adjusted *P*-values < 0.05 are shown in blue and the 20 proteins with the highest absolute log_2_ fold change are labelled. *P*-values were computed by limma-based linear models with Bonferroni’s correction for multiple testing. **C:** Metascape(Zhou *et al*, 2019) enrichment heatmap showing functional clusters of phosphoproteins affected by *S. aureus* exposure. **E:** Metascape(Zhou *et al*, 2019) bar graph showing top non-redundant functional clusters of phosphoproteins enriched in the vehicle control/*S. aureus* condition versus C5a/*S. aureus* condition.

Our dataset suggests that C5a exposure that precedes pathogen encounter prevents effective signalling through the phagosomal maturation pathways, and links intracellular signalling to the prolonged functional impairment noted in this context. The other major cluster of differentially phosphorylated proteins were nuclear and nuclear membrane proteins, many of which are involved in mitosis and nuclear envelope integrity.

### C5a induces an impairment in phagosomal acidification, distinct from the impairment in ingestion

The phosphoproteomic signature of altered phagosomal maturation following C5a exposure, and the involvement of V-ATPase suggested that C5a had effects beyond impaired ingestion of bacteria. To disentangle the effects of phagocytic ingestion and phagolysosomal acidification, *S. aureus* bioparticles co-labelled with the pH-insensitive dye AF488 and pHrodo™ red were used. Neutrophils ingested particles, and then subsequently acidified the phagosome, a process which could be ablated by the addition of the V-ATPase inhibitor bafilomycin (Bowman *et al*, 1988) (Figures 5A and B). C5a pre-treatment increased the proportion of neutrophils that failed to ingest particles (Figure 5C) and increased the population that ingested particles but failed to acidify the phagosome (Figure 5D). Recent reports suggest that C5a induces Na^+^/H^+^ exchanger-1 (NHE-1)-mediated cytoplasmic alkalinisation (Denk *et al*, 2017a). An NHE-1 inhibitor did not alter the C5a-mediated effect on phagosomal acidification (Figure 5E), suggesting that the pathways mediating these two effects of C5a on neutrophils are distinct. Furthermore, we confirmed previous work (Huber-Lang *et al*, 2002b) showing C5a impaired ROS production (Figure S8), which in combination with the current findings, suggests C5a induces a generalised failure of phagosomal maturation in addition to its effect on phagocytic ingestion.

**Figure 5:**
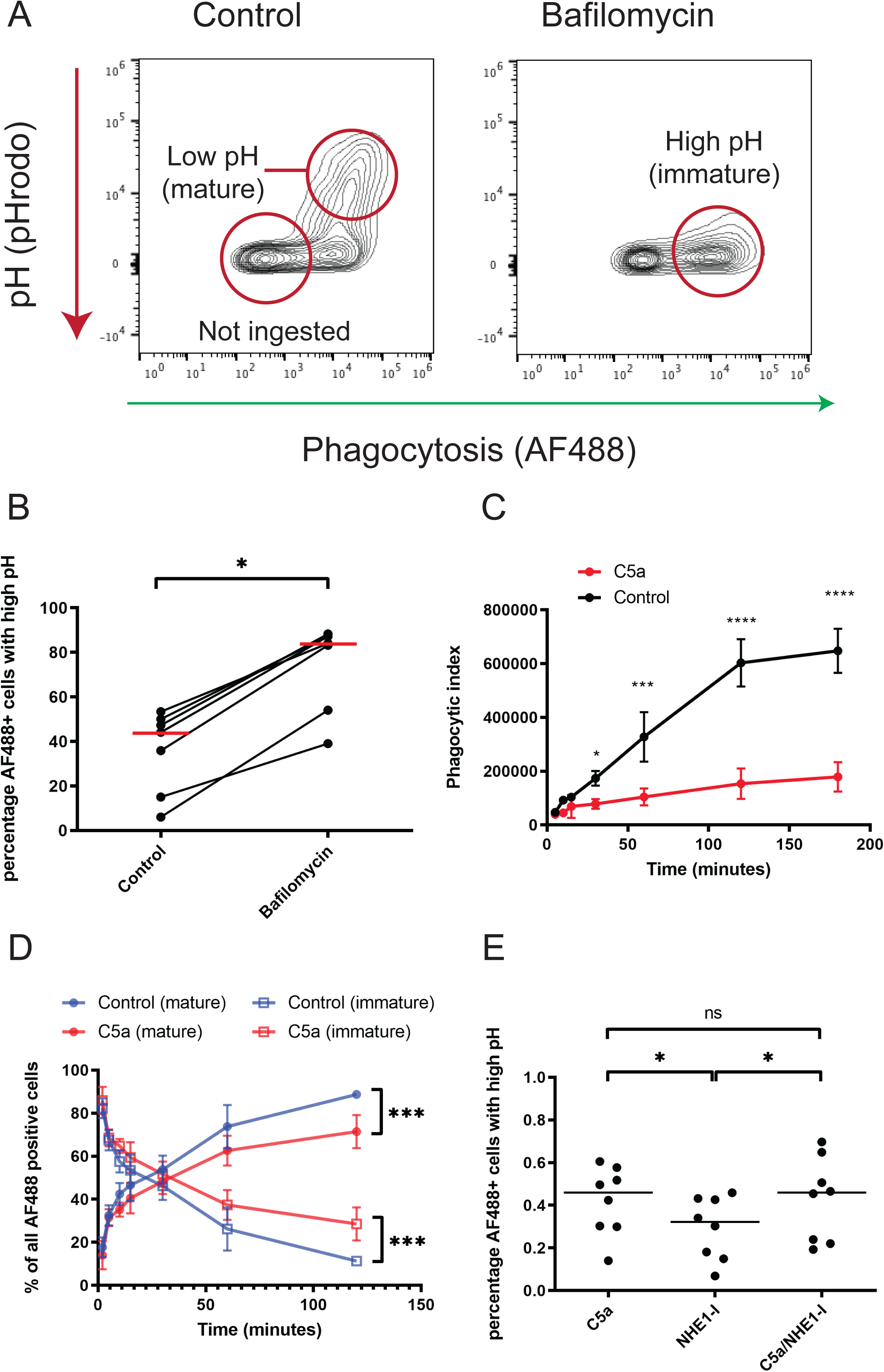
C5a induces an impairment in phagosomal acidification, distinct from the impairment in ingestion. **A:** Exemplar flow cytometry plots of whole blood pre-treated with vehicle control or bafilomycin A (60 min; 100nM) prior to exposure to 5 µg/mL co-labelled AF488/pHrodo red *S. aureus* for 120 min. Both phagocytosis (x-axis) and phagosomal pH (y-axis) can be measured simultaneously in the same population of cells. pHrodo™ fluorescence increases with decreasing pH, indicating phagosomal maturity as shown. **B:** Conditions as in A. Data are shown as individual data points with mean for n=7 individual donors. *P* = 0.016 by Wilcoxon’s test. **C:** Whole blood was pre-treated with vehicle control or C5a (300 nM; 60 minutes) prior to exposure to phagocytosis probe for 180 min. Phagocytosis without maturation (i.e. AF488 signal) is shown. Data are shown as mean and SD of n = 5 individual donors. \*\*\*\**P* < 0.0001 by repeated-measures two-way ANOVA with Bonferroni’s multiple comparisons test. **D:** Conditions as in C. The percentage of *S. aureus* particle positive (AF488+) cells with low pH (mature) and high pH (immature) phagosomes is shown for control and C5a-treated conditions. Data are shown as mean and SD of n = 5 individual donors. \*\*\**P* < 0.001 by repeated-measures two-way ANOVA with Bonferroni’s multiple comparisons test. **E**: Whole blood was pre-treated with C5a, NHE-1 inhibitor (5μM), or both, then exposed to maturation probe for 60 min. The percentage of AF488+ cells with high pH (immature) phagolysosomes is shown. Data are shown as individual data points with median from n = 7 individual donors. *P* = 0.0080 by Friedman’s test, \**P* < 0.05 for Dunn’s test of multiple comparisons, ns = non-significant.

### VPS34 inhibition impairs phagosomal acidification

The differential phosphoprotein analysis (Table 1) and phagosomal acidification assays (Figure 5) demonstrated impaired phagosomal maturation after exposure to C5a. As noted, several of the phosphoproteins that were differentially phosphorylated are known interactors with PI3P. The phosphatidylinositol 3-kinase VPS34 is the dominant source of PI3P in mammalian cells (Devereaux *et al*, 2013). Although VPS34 itself was detected, its phosphorylation status was not significantly altered. However, the finding that C5a altered the phosphorylation status of PI3P-responsive proteins led us to explore the role of VPS34 in phagosomal acidification. We used the selective inhibitor, VPS34IN1 (Bago *et al*, 2014) to examine the role of this enzyme in phagosomal acidification, and how this related to the defect induced by C5a. VPS34IN1 did not alter the percentage of neutrophils that underwent phagocytosis (Figure 6A and time-course in E) but did lead to a reduction in the overall number of particles ingested (Figure 6B) and a more marked reduction in pHrodo signal (Figure 6C and time course in F), indicating VPS34IN1 impairs phagosomal acidification. VPS34 inhibition also led to an impairment in the killing of *S. aureus* (Figure 6D), similar to that observed with C5a (Figure S1D) without a significant reduction in phagosomal ROS production (Figure S9).

**Figure 6:**
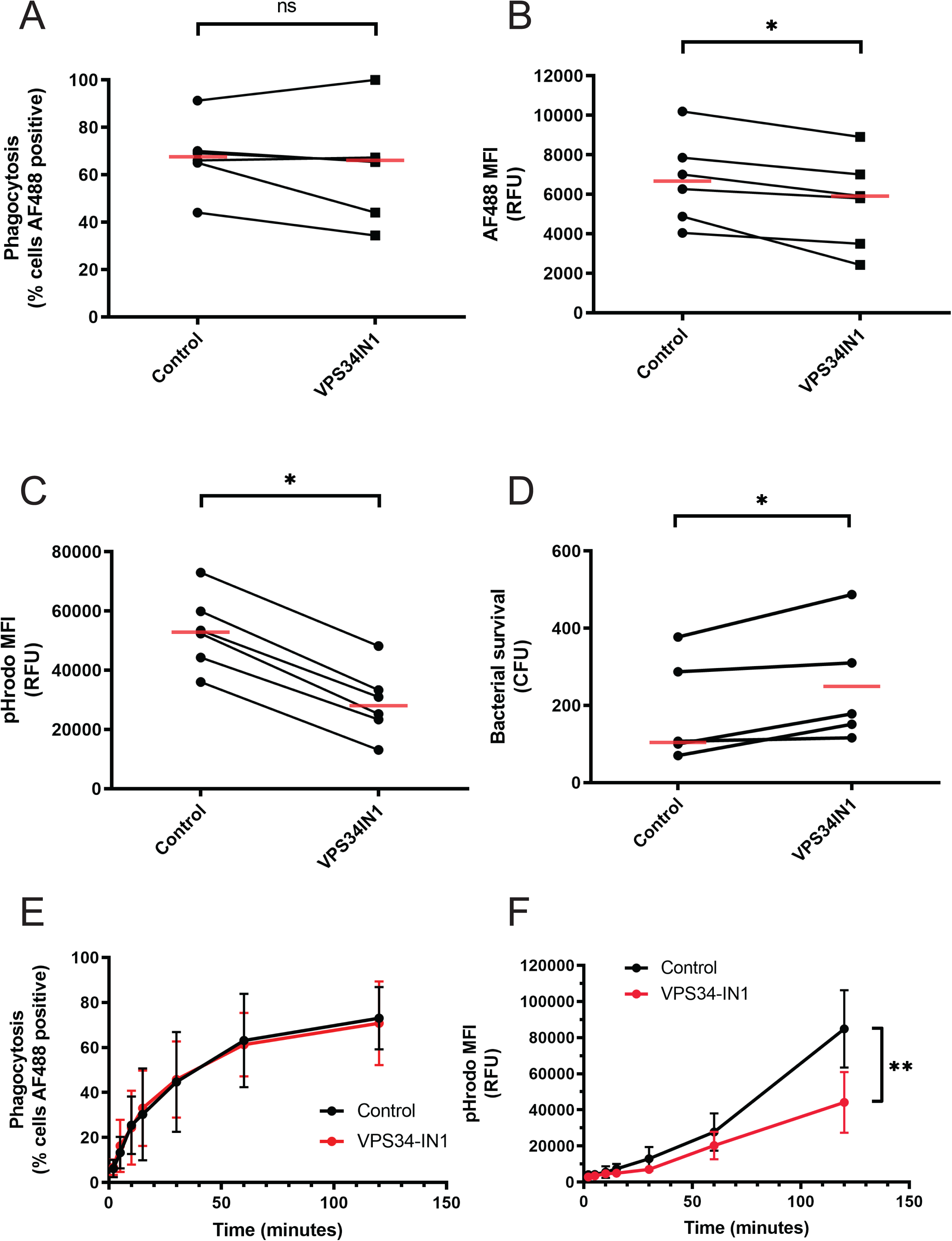
VPS34 inhibition impairs phagosomal acidification. Whole blood was pre-treated with vehicle control or VPS34IN1 (1 µM; 60 min) prior to addition of 5 µg/mL maturation probe (A-D), or live *S. aureus* (E), for 120 minutes prior to analysis. **A**: Percentage of neutrophils that have phagocytosed bioparticles. *P* = 0.31 by Wilcoxon’s test. n = 6 individual donors. **B**: MFI of ingested particles, indicating relative quantity of phagocytosis. *P* = 0.03. by Wilcoxon’s test. n=6 individual donors. **C**: pHrodo™ Median Fluorescent Intensity (MFI), indicating phagosomal acidification. *P* = 0.03. by Wilcoxon’s test. n=6 individual donors. **D**: After phagocytosis of live bacteria, human cells were lysed in alkaline dH_2_O and surviving bacteria were incubated overnight on blood agar. Bacterial survival was quantified by counting colonies. *P* = 0.03 by paired t-test, n=5 individual donors. **E-F:** Whole blood was processed as above with quantification of phagocytosis (E) and acidification (F) at the indicated time points. There was a reduction in phagosomal acidification as shown but no change in percentage of cells that underwent phagocytosis. Data are shown as mean and SD of n=5 individual donors. \*\**P* = 0.0058 for drug treatment by repeated measures two-way ANOVA with Bonferroni’s multiple comparisons test.

### Neutrophils from critically ill patients exhibit defective phagosomal acidification

To establish the relevance of our findings to the clinical setting, we used our assay of phagosomal acidification to interrogate neutrophils obtained from critically ill patients and healthy volunteers. We assessed neutrophil function in critically ill patients, defining neutrophil dysfunction as phagocytosis of <50% in our previously established zymosan assay (Figure 7A), a threshold associated with a markedly increased risk of nosocomial infection (Pinder *et al*, 2018; Conway Morris *et al*, 2009, 2011). Using our phagosomal acidification assay, we then compared patients with dysfunctional neutrophils to critically ill patients with functional neutrophils and healthy controls. Dysfunctional neutrophils exhibited a failure of phagosomal acidification (Figure 7B) that was not seen in patients with functional neutrophils. Furthermore, we observed a correlation between C5aR1 expression (decreased after C5a exposure) and phagocytosis (Figure 7C) and an inverse correlation between C5aR1 expression and phagosomal acidification (Figure 7D), though the latter correlation did not reach statistical significance. The patients with dysfunctional and functional neutrophils could not be readily identified by clinical factors such as severity of illness or precipitating insult (Table S2). These data provide evidence of dysfunctional phagosomal acidification in critically ill patients and imply a role for C5a in driving this dysfunction.

**Figure 7:**
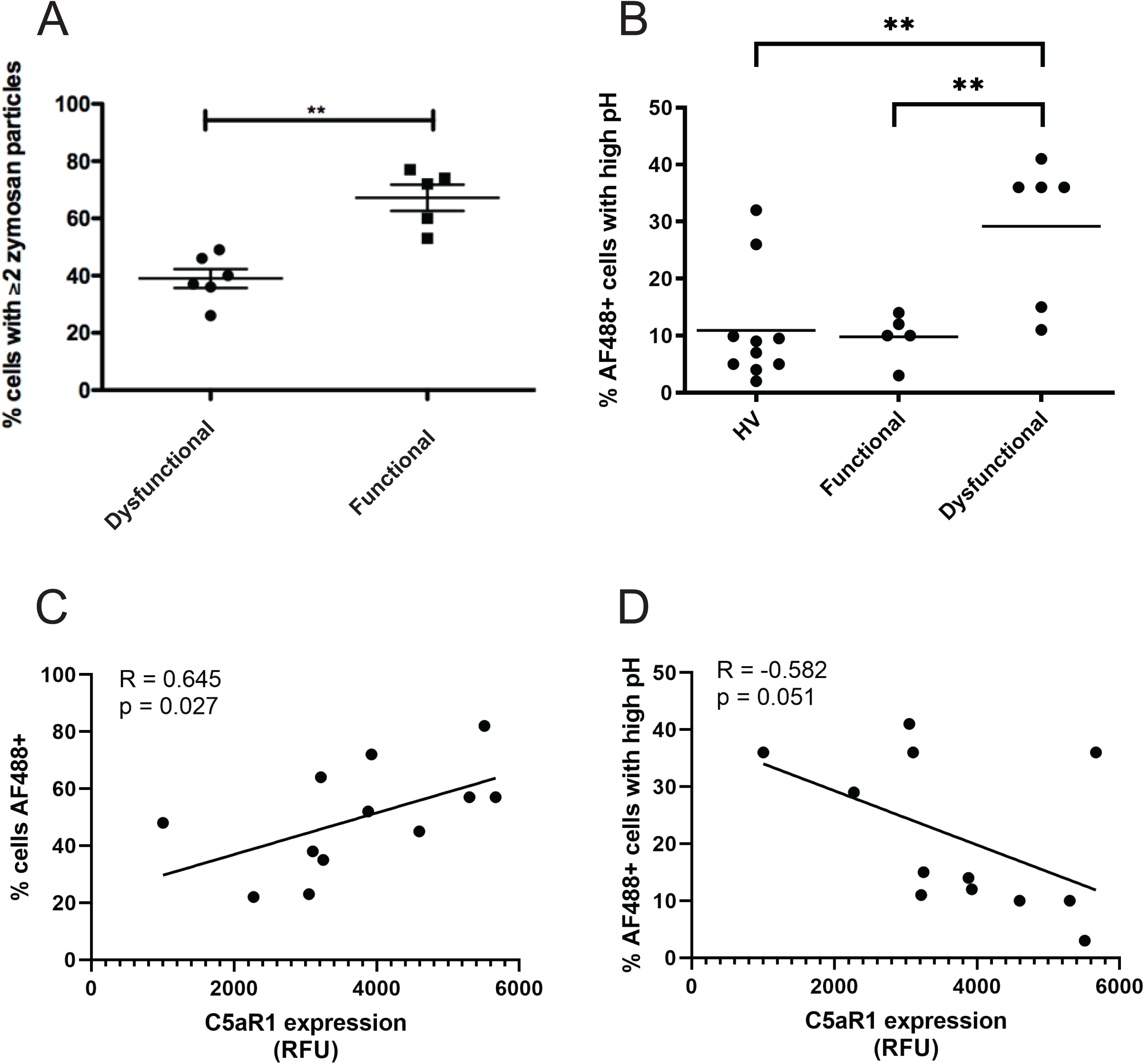
Neutrophils from critically ill patients exhibit defective phagosomal acidification. **A**: Zymosan-based assay demonstrating differentially impaired phagocytosis in critically ill patients. Data are shown as individual patients/controls with median values indicated. n = 6 patients with dysfunctional neutrophils and 5 patients with functional neutrophils respectively. ** *P*=0.004 by Mann-Whitney U-test. **B**: Neutrophil phagosomal acidification was assessed in whole blood from critically ill patients using the maturation probe. Patients were classed as dysfunctional using the assay from A. Data are shown as individual patients/controls with mean from n = 6 patients with dysfunctional neutrophils, 5 patients with functional neutrophils and 10 healthy controls respectively. *P* = 0.04 by one-way ANOVA. \*\**P* < 0.01 by Holm-Sidak’s test of multiple comparisons. **C, D**: C5aR1 expression was assessed by flow cytometry and correlated (Spearman) with phagocytosis (C) and phagosomal acidification (D) for n = 12 patients. NB: One patient’s cells did not adhere to tissue culture plastic for the zymosan assay, thus they could not be assigned to dysfunctional or non-dysfunctional groups shown in A and B. C5aR1 expression and maturation probe data was available to allow inclusion in correlation analyses in C and D, hence the difference in numbers between these figures.

## Discussion

Our data demonstrate that C5a induces both a prolonged defect in phagocytosis of relevant pathogens (*S. aureus* and *E.coli*), and persistent signalling across multiple pathways for some hours after the well characterised initial signalling events such as ionised calcium flux (Blackwood *et al*, 1996) and PIP3 generation (Houslay *et al*, 2016). This finding supports the proposal that persistent C5a-induced signalling may mediate the neutrophil dysfunction observed in critically ill patients (Conway Morris *et al*, 2009, 2011).

To our knowledge, the data presented here (Figures 3-5) represent the deepest sequencing of the human neutrophil proteome and phosphoproteome (Muschter *et al*, 2015; Tak *et al*, 2017; McLeish *et al*, 2013). These data provide a phosphoproteomic assessment of the human neutrophil response to *S. aureus* and C5a. Unlike transcriptomic data (Juss *et al*, 2016; Rorvig *et al*, 2013; Kobayashi *et al*, 2002), phosphoproteomics provides a direct assessment of mediators that are likely to have functional implications, especially in short-lived cells such as neutrophils (Luerman *et al*, 2010; Fessler *et al*, 2002) and early pathogen exposure timepoints, as examined in this study.

The marked changes observed in phosphoproteins in response to *S. aureus* are perhaps unsurprising, as the response to and clearance of bacteria are primary functions of neutrophils. Many of the pathways identified (Figures 4 and S4) are consistent with established literature on neutrophil responses to *S. aureus*, and indeed other bacteria, including activation of PI3K (Li *et al*, 2016), toll-like receptor signalling (Jann *et al*, 2011) and neutrophil degranulation (McGovern *et al*, 2011).

The enrichment of PI3K and Rho GTPase signalling on C5a stimulation are in keeping with our previous identification of key roles for these molecules in C5a-mediated functional deficits in neutrophils (Conway Morris *et al*, 2009, 2011; Scott *et al*, 2015). The marked suppression of the phosphorylation response to *S.aureus* induced by C5a pre-treatment is not simply a response to reduced particle ingestion. Fifteen minutes after pathogen contact there were limited differences in the ingestion rates between C5a and control treatments, and these became more marked over time (Figure 1). Furthermore, the differential analysis of C5a/*S. aureus* versus vehicle control/*S. aureus* conditions identified defects in specific signalling pathways, most notably those involving endosomal trafficking. This led us to examine the process of phagosomal maturation, and to identification of a C5a-induced failure of phagosomal acidification (Figure 5) with similar findings in critically ill patients (Figure 7). Failure of phagosomal maturation and intracellular killing has been described in primary immune deficiency (Buvelot *et al*, 2017), but has not previously been described as part of the immuno-paresis of critical illness. Impaired phosphorylation in pathways involving nuclear envelope breakdown and nuclear pore disassembly by C5a was unanticipated. The functional relevance of these changes remains unclear, though they may be early processes in the formation of non-lethal DNA-containing neutrophil extracellular traps (NETs) (Pilsczek *et al*, 2010).

Important signalling proteins involved in the process of phagosomal maturation (such as RAB7A, TOM1 and ZFYVE16) can be recruited to the phagosomal membrane by PI3P produced predominantly by VPS34 (Botelho *et al*, 2000; Levin *et al*, 2016; Sorkin & Von Zastrow, 2009). Both ZFYVE16 and TOM1 phosphorylation were impaired by C5a exposure. We investigated the role of VPS34 as a mediator of neutrophil bactericidal function, and found that selective VPS34 inhibition produced a similar impairment in phagosomal acidification to that observed with C5a (Figure 6). The finding that a similar defect could be induced by inhibiting VPS34, the dominant source of PI3P in neutrophils (Devereaux *et al*, 2013), adds further validation to the pathway signature identified in the phosphoproteomic profile.

Ellson and colleagues (Ellson *et al*, 2001) demonstrated that PI3P plays an important role in targeting neutrophil oxidase components to phagosomal membranes and its importance in phagosomal maturation has also been identified in *Dictyostelium discoideum* (Buckley *et al*, 2019), murine macrophages, and macrophage-like cell lines (Naufer *et al*, 2018). However, the role of VPS34 in human neutrophils has previously been inferred indirectly (Anderson *et al*, 2008), owing to prior lack of selective inhibitors and the difficulties of genetically manipulating human neutrophils. Anderson and colleagues (Anderson *et al*, 2008) demonstrated a role for VPS34 in NADPH oxidase-mediated reactive oxygen species generation in neutrophils. We found a non-significant reduction in ROS production (Figure S9) that was much less marked than the effect on phagosomal acidification. The reasons for these divergent findings are uncertain, though may include differences in ROS measurement assays, our use of a selective VPS34 inhibitor, and differences between primary human neutrophils and cell lines. The mechanism by which VPS34 inhibition impairs killing of *S. aureus* requires further investigation, as phagosomal acidification is not thought to be critical to this process (Lacoma *et al*, 2017) and it is likely that the enzyme inhibition leads to further defects in phagosomal maturation. It is intriguing to note that whilst VPS34 inhibition does not reduce the percentage of cells that undergo phagocytosis (Figure 6A), consistent with previous work (Anderson *et al*, 2008), it does reduce the number of particles ingested (Figure 6B). This is consistent with the finding that ingestion of an initial target facilitates ingestion of a subsequent target (Figure S2), implying a capacitive stage of phagocytosis. That VPS34 inhibition impairs this capacitive phase suggests a hitherto undescribed relationship between phagosomal maturation and the capacity of cells to ingest particles.

Our data also demonstrate that the timing of C5a exposure (before, alongside, or after pathogen encounter) has an important effect on neutrophil function. Only pre-exposure to C5a impaired subsequent neutrophil phagocytosis (Figure 2). Reduced C5aR1 availability for ligation by C5a is unlikely to explain this observation, as C5aR1 downregulation is induced by multiple agents that do not have the same effect on phagocytosis (Figure S1D). The pathways enriched under C5a conditions are also represented amongst those enriched for following *S. aureus* exposure, with all but 8 of the C5a-induced significantly altered phosphoproteins also being identified following *S. aureus.* It is therefore plausible that signalling induced by *S. aureus* overwhelms C5a-induced phosphorylation events unless they were established prior to *S. aureus* exposure. Indeed, the signalling induced by initial ingestion may facilitate subsequent ingestion, as identified in the sequential exposure assay (Figure S2), actively mitigating against C5a-induced suppression. We did not identify a dose at which C5a enhanced phagocytosis in this work (Figure S1A) or indeed in our previous work (Conway Morris *et al*, 2009, 2011). Collectively, these findings imply that the divergent findings on the effect of C5a on phagocytosis when C5a is added exogenously, as in this report or generated by direct addition of bacteria to blood as reported by Mollnes and colleagues (Mollnes *et al*, 2002; Brekke *et al*, 2007), are due to the temporal relationship between exposures rather than a biphasic dose-response relationship.

This finding of distinct temporal responses to C5a suggests that where neutrophils encounter bacteria and C5a at the same time, such as at the site of infection, the phagocytic response is not impaired. When complement activation spills over systemically and C5a exposure precedes neutrophil-bacterial interactions - as occurs with systematic inflammation in critical illness and sepsis - dysfunction occurs, impairing the host’s ability to respond to invasive infections or subsequent bacterial insults (Conway Morris 2013, 2018).

This study was conducted entirely in primary human neutrophils, using C5a, an established, clinically relevant modulator of neutrophil function that has been linked to a range of adverse outcomes in critically ill patients. The use of clinically relevant pathogens, and the development of a whole-blood bacteraemia model, increases the relevance of our study to the *in-vivo* situation. Impaired ingestion of zymosan by patient neutrophils has been associated with adverse outcomes including development of subsequent nosocomial infection (Conway Morris *et al*, 2011). The finding that patients with such impairment also manifest impaired phagosomal acidification that correlates with markers of C5a exposure (Figure 7) suggests that the identified mechanisms may be clinically relevant.

Several potential limitations should be highlighted. The phosphoproteomic response to *S. aureus* was evoked with heat-killed bacterial particles, conjugated with fluorescent dyes, and these may not fully reflect the response to live bacteria, although they do allow parallel functional assessment and standardisation of the stimulus between donors and across research sites. Although whole blood is a more physiologically relevant than cell-culture media, it remains an abstraction from the situation *in-vivo*, as it must be anticoagulated and does not involve normal flow or interaction with a vascular endothelium. Furthermore, the model may not reflect the function of neutrophils that have migrated into tissues, where most bacterial infections occur. Technical limitations currently prevent efficient phosphoproteomic assessment of cells from whole blood, and therefore isolated cells with the inherent *in-vitro* artefacts must be used.

In conclusion, we have demonstrated the role of C5a in mediating neutrophil dysfunction in the clinically relevant setting of *S. aureus* and *E. coli* bacteraemia, and demonstrated that the effects of C5a can persist for many hours and which is dependent on the temporal sequence of C5a and bacterial exposure. We also describe the neutrophil phosphoproteomic response *to S. aureus*, and to prolonged exposure to C5a. This approach identified a defective phagosomal maturation signature induced by C5a, likely involving modulation of Class III PI3K-dependent pathways. Further, we have shown the functional manifestation of this phosphorylation signature in a model of bacteraemia. Finally, the clinical relevance of this failure of phagosomal acidification was observed in critically ill patients. A deeper understanding of the biology of neutrophil dysfunction in critical illness is key to developing effective treatments for a phenomenon associated with multiple adverse clinical outcomes.

## Materials and methods

### Donors

Ethical permission for obtaining peripheral venous blood from healthy volunteers was provided by the Cambridge Local Research Ethics Committee (REC reference 06/Q0108/281) and all donors provided written, informed consent. Critically ill patient blood samples were obtained under an approval granted by the North East-Newcastle & North Tyneside 2 Research Ethics Committee (REC reference: 18/NE/0036). Inclusion and exclusion criteria are detailed in the supplemental methods. Assent was provided by a personal or nominated consultee.

Further details of methods and reagents described below are available in the supplementary materials.

### Neutrophil isolation

Neutrophils were isolated from citrated peripheral venous blood by using a modification of the discontinuous plasma-Percoll density gradient centrifugation technique initially described by Böyum in 1968.(Boyum, 1968)

### Phagocytosis of pHrodo™ S. aureus and E. coli Bioparticles by purified neutrophils

Purified human neutrophils, suspended in Iscoves Modified Dulbecco’s Medium (IMDM) with 1 % autologous serum at a concentration of 5 × 10^6^/mL, were incubated in microcentrifuge tubes with purified human C5a or vehicle control. pHrodo-conjugated *S. aureus* or *E. coli* bioparticles were opsonised, in 50 % autologous serum for 30 min prior to be being added to the suspended cells. Analysis was by flow cytometry (Attune NxT, Thermofisher)

### No-wash, no-lyse whole blood assay of neutrophil phagocytosis and ROS production

Blood, collected into argatroban 150 µg/mL, was treated with inhibitors or priming agents as indicated in the respective figure legends, before being exposed to *S. aureus* pHrodo™/dihydrorhodamine (DHR) or *E.coli* pHrodo™. Aliquots were stained on ice with anti-CD16 antibody, diluted and analysed by flow cytometry (Attune NxT).

In variations on this assay, *S. aureus* particles labelled with the pH-insensitive dye AlexaFluor (AF)488 or dual labelled with AF488 and pHrodo red were used. pHrodo red conjugation of AF488 *S. aureus* was performed in-house using the pHrodo particle labelling kit (Thermofisher). Fluorescence of extracellular particles was quenched with trypan blue (0.1mg/mlL).

Patient samples were analysed in a different laboratory that did not have access to an Attune Nxt flow cytometer, to fit with established workflows in this laboratory red cells were lysed using Pharmlyse (BD Bioscience, Wokingham, UK) followed by washing twice using a Facswash Assistant (BD Bioscience) prior to undertaking flow cytometry (Fortessa, BD Bioscience).

### Bacterial killing assay – whole blood

Methicillin-sensitive *S. aureus* (MSSA) bacteria (strain ASASM6, kind gift from Prof Gordon Dougan, University of Cambridge) were grown to early log-phase. Blood was collected into argatroban and incubated with bacteria for 1 hour. Human cells were lysed by addition of pH 11 distilled water for 3 minutes before plating of serial dilutions on Colombia blood agar.

### Preparation of whole human neutrophil lysates for phosphoproteomics

Neutrophils were isolated from whole blood as detailed above, and resuspended in RPMI 1640 media containing 10 mM HEPES with 1 % autologous serum (AS) at a concentration of 1×10^7^ cells/mL.

### Proteomic and phosphoproteomic studies

Triplicates of 1×10^7^ neutrophils were treated with vehicle control or C5a (100 nM, 60 minutes) at 37 °C before addition of pHrodo™ *S. aureus* (15 µg/mL). Phagocytosis was allowed to occur for 15 minutes. Aliquots were withdrawn from each triplicate and pooled at the indicated timepoints. Cells were centrifuged at 400 g for 5 min at 4 °C, supernatants aspirated, and cell pellets snap frozen in liquid nitrogen. Cells were lysed by the addition of 0.5 % sodium dodecyl sulphate (SDS)/0.1 M triethylammonium bicarbonate (TEAB) buffer and sonication, before undergoing centrifugation, trypsin digestion, tandem mass tag labelling, fractionation, phosphopeptide enrichment, and liquid chromatography and tandem mass spectrometry (LC-MS/MS) analysis. The experimental schematic can be seen in Supplemental Figure S10.

### Statistical analysis of wet laboratory data

Data are presented as individual data points with summary statistics (median and interquartile range (IQR) or mean and standard deviation (SD) according to whether data are normally distributed. Parametric or non-parametric statistical tests were applied as appropriate after data were tested for normality using the D’Agostino-Pearson test. Tests used for comparisons are indicated in figure legends. Two-tailed *P* values were computed, *P* < 0.05 was considered statistically significant. Non-significant differences have not been indicated in figures for clarity. Statistical analyses were undertaken using GraphPad Prism v8.0 (GraphPad Software; San Diego; California).

### Statistical analysis of phosphoproteomics data

Spectral .raw files from data dependent acquisition were processed with the SequestHT search engine on Thermo Scientific Proteome Discoverer™ 2.1 software. Data were searched against both human and *S. aureus* UniProt reviewed databases at a 1 % spectrum level false discovery rate (FDR) criteria using Percolator (University of Washington). MS1 mass tolerance was constrained to 20 ppm, and the fragment ion mass tolerance was set to 0.5 Da. TMT tags on lysine residues and peptide N termini (+229.163 Da) and methylthio (+45.988 Da) of cysteine residues (+45.021 Da) were set as static modifications, while oxidation of methionine residues (+15.995 Da) and deamidation (+0.984 Da) of asparagine and glutamine residues were set as variable modifications. For TMT-based reporter ion quantitation, we extracted the signal-to-noise (S:N) ratio for each TMT channel. Parsimony principle was applied for protein grouping.

Peptide and phosphopeptide intensities were normalised across conditions using median scaling and then summed to generate protein and phosphoprotein intensities. Proteins and phosphoproteins were independently identified and quantified in all samples from all four donors; species not meeting these criteria were excluded from subsequent analysis. Log base 2 fold change (Log2FC) was calculated between conditions of interest, compared across n = 4 donors and tested for statistical significance by limma-based linear models with Bonferroni’s correction for multiple testing. Hierarchical clustering using Euclidean distance was undertaken on the entire dataset. Heatmaps and volcano plots were generated as shown in Results. Statistical analyses were performed in RStudio (RStudio Team, 2016) using the qPLEXanalyzer (Papachristou *et al*, 2018) package, and plots were produced using the ggplot2 (Wickham, 2016) package.

### Data sharing statement

The mass spectrometry proteomics data have been deposited to the ProteomeXchange Consortium via the PRIDE (Perez-Riverol Y *et al*, 2019) partner repository with the dataset identifier PXD017092 and will be made public on acceptance after peer-review

## Supporting information

Supplemental results and methods

## Acknowledgements

We gratefully acknowledge the generous gift of live methicillin sensitive *S. aureus* from Prof Gordon Dougan, University of Cambridge. This research was supported by the Cambridge NIHR BRC Cell Phenotyping Hub and the Cambridge NIHR Biomedical Research Centre. In particular we wish to thank Esther Perez and Natalia Savinykh for their advice on flow cytometry.

The views in this manuscript represent the views of the authors alone, and not those of the National Institute for Health Research or the Department of Health and Social Care.

## Authorship contributions (CRediT)

AJTW: Conceptualisation, formal analysis, investigation, methodology, validation, visualisation, writing – original draft, review and editing.

AV: Investigation, methodology, writing - review and editing.

MHRS: Investigation, methodology, writing - review and editing.

JS: Investigation, methodology, writing - review and editing.

CZ: Investigation, writing - review and editing.

CGT: Investigation, methodology, writing - review and editing.

KK: Methodology, data curation, software, formal analysis, writing - review and editing.

CSDS: Project administration, resources, supervision, writing - review and editing.

AJS: Project administration, resources, supervision, writing - review and editing.

DKM: Project administration, resources, supervision, writing - review and editing.

CS: Methodology, validation, project administration, resources, supervision, writing - review and editing.

ERC: Conceptualisation, formal analysis, funding acquisition, methodology, project administration, resources, supervision, writing - review and editing.

KO: Project administration, resources, supervision, writing - review and editing.

ACM: Conceptualisation, formal analysis, funding acquisition, investigation, methodology, project administration, resources, supervision, validation, writing – original draft, review and editing.

