## Supplemental results and methods for "Complement C5a impairs phagosomal maturation in the neutrophil through phosphoproteomic remodelling"

Supplemental Table 1: Marked phosphoproteome changes with treatment

This table shows statistically significant numbers of protein/phosphoprotein alterations in the total proteome and total phosphoproteome between the conditions indicated. These alterations are shown as both the absolute number and as a percentage of the respective proteome/phosphoproteome.

|  | Total proteome (4859 proteins quantified) | | Total phosphoproteome (2712 phosphoproteins quantified) | |
| --- | --- | --- | --- | --- |
| Conditions compared | Number of proteins significantly altered | Percentage of total proteome significantly altered | Number of phosphoproteins significantly altered | Percentage of total phosphoproteome significantly altered |
| Ctrl-untreated | 0 | 0.0% | 16 | 0.6% |
| C5a-ctrl | 7 | 0.1% | 111 | 4.1% |
| *S. aureus*-ctrl | 95 | 2.0% | 858 | 31.6% |

| Parameter | Patients with  Dysfunctional neutrophils (n=6) | Patients with non-dysfunctional neutrophils  (n=5) |
| --- | --- | --- |
| Median (range) age | 62 (44-77) | 65 (48-81) |
| % female (n) | 33% (2) | 20% (1) |
| Median (Interquartile range) admission APACHE II score | 19.5 (17.25-29) | 16.5 (12-21) |
| % with suspected sepsis on admission  (of which culture positive) | 66% (4)  100% (4) | 100% (5)  60% (3) |

Supplemental Table 2: Clinical characteristics of critically ill patients who underwent neutrophil functional assessment

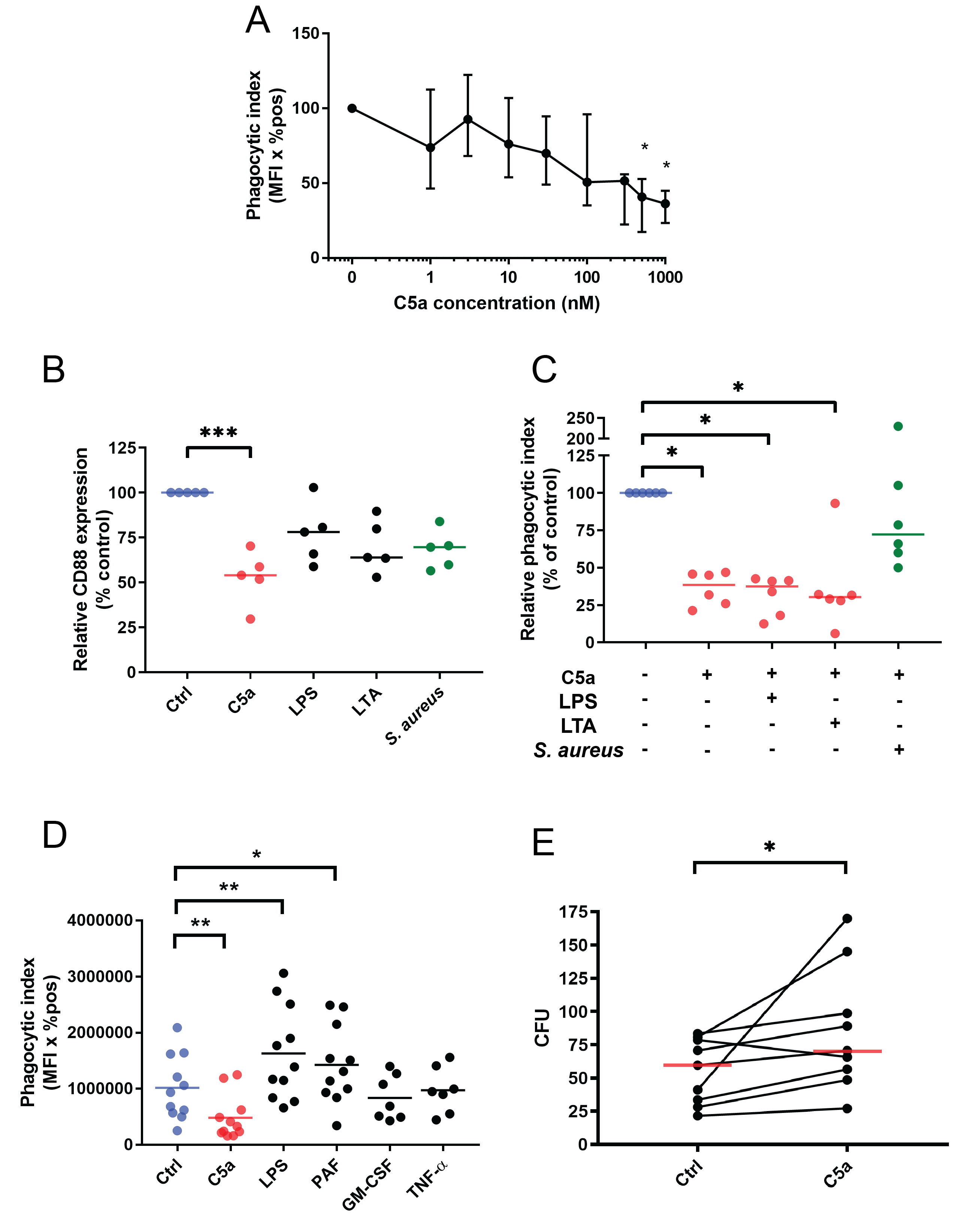

Supplemental figure 1: C5aR1 changes and phagocytosis with inflammatory stimuli, neutrophil killing of *S. aureus*

**A:** Whole blood was pre-treated with the indicated concentrations of C5a for 60 minutes prior to phagocytosis of pHrodo bioparticles. Data are shown as the median and IQR of 4 independent experiments relative to control-treated blood. Friedman p-value = 0.0024, *p < 0.05 by Dunn’s multiple comparisons.

**B**: Whole blood was incubated with the indicated inflammatory stimuli for 60 minutes prior to measurement of neutrophil C5aR1 expression by flow cytometry: 300 nM C5a; 100 ng/mL LPS; 10 µg/mL LTA; *S. aureus* bioparticles 10 µg/mL. Individual data points and their median values from n = 5 independent experiments are shown. Friedman’s test with p-value = 0.0003, ***p < 0.001 by Dunn’s multiple comparisons test.

**C**: Whole blood was pre-incubated with control or inflammatory stimuli as in (A) and then with 300 nM C5a for 60 minutes. Cells were then allowed to phagocytose 10 µg/mL *S. aureus* pHrodo™ red Bioparticles for 60 minutes before phagocytosis was assessed by flow cytometry as previously described. Individual data points and their median values from n = 6 independent experiments are shown. Friedman’s test with p-value = 0.0038, *p < 0.05 by Dunn’s multiple comparisons test

**D**: Whole blood neutrophil phagocytosis after pre-treatment with various agonists. Concentrations and pre-treatment durations were as follows: Ctrl (PBS) 60 minutes; C5a 300 nM, 60 minutes; LPS 100 ng/mL, 60 minutes; PAF 1 μM, 5 minutes; GM-CSF 10 ng/mL, 30 minutes; TNF-α 20 ng/mL, 30 minutes. ANOVA p-value < 0.0001 for both D and E, *p < 0.05, **p < 0.01 for Dunnett’s multiple comparisons. Data are shown as individual data points with means for n = 11 (E) and n = 7 (F) individual donors.

**E**: Purified human neutrophils were pre-treated with 100 n C5a for 60 minutes prior to incubation with live *S. aureus* (strain ASASM6) for 60 minutes at an MOI of 10:1 bacteria per neutrophil. Neutrophils were lysed, bacteria plated and grown overnight and CFUs quantified by visual inspection. Individual data points from n = 9 independent experiments are shown. C5a pre-treatment resulted in an increase in median CFU count of 30.8 % relative to control. p = 0.02 by Wilcoxon.

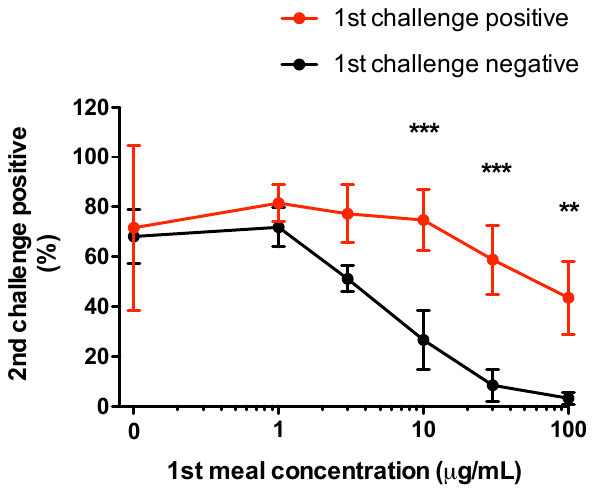

B

1st challenge

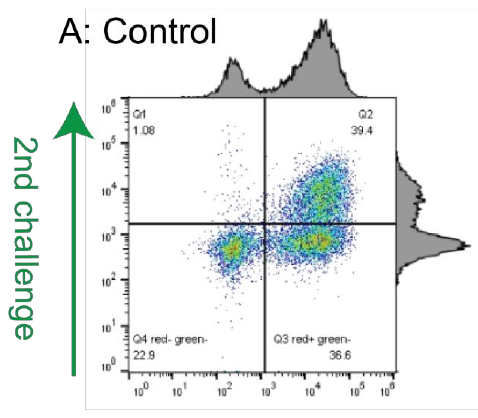

C

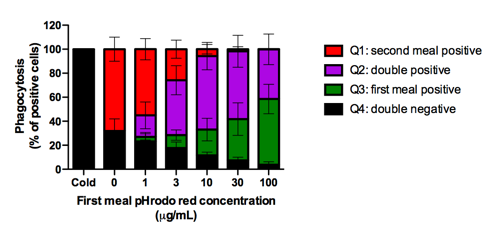

**Supplemental figure 2: whole blood sequential ‘feeding’ assay indicates a facilitatory relationship between prior and subsequent ingestion of phagocytic targets.**

**A:** Example flow plot showing results from healthy donor blood exposed to 15µg/ml of pHrodo red *S. aureus* for 30 minutes followed 15µg/ml of pHrodo green *S. aureus* for 30 minutes. Cells divide into 3 populations, bottom left quadrant low/no phagocytosis, bottom right, single phagocytosis, top right double phagocytosis. Of note very few cells ingest the second meal having not ingested the first (top left quadrant).

**B:** Increasing the concentration of the first ‘meal’ reduces the proportion of cells eating a standard (15 µg/mL) sized second ‘meal’, indicating a fundamental limit on the capacity of neutrophils to ingest particles. Increasing the size of the first meal leads to progressive selection of a resolutely non-phagocytic population corresponding to Q1 shown in panel A. Data from n=5 individual donors, P<0.0001by repeated measures two way ANOVA. *** p<0.001, **P<0.01 by Bonferroni’s post-hoc test.

**C:** Proportions of cells in each quadrant with increasing size of first ‘meal’. Data from n=5 individual donors

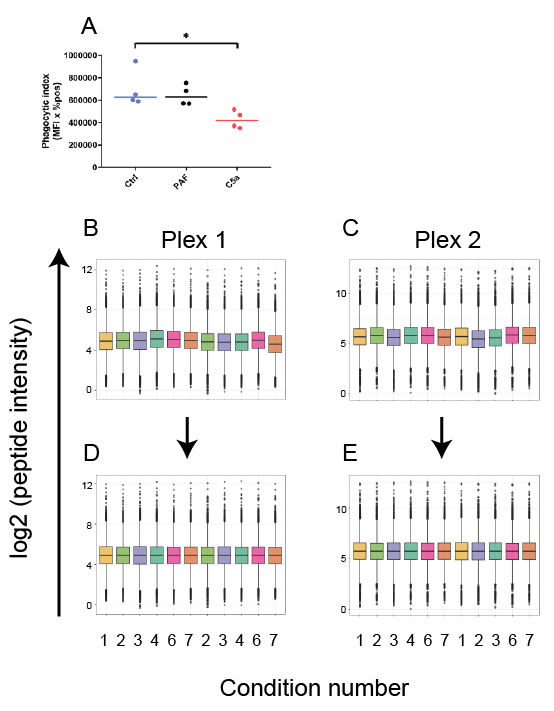

Supplemental figure 3: technical validation of phosphoproteome assessment

**A**: Neutrophils were pre-treated with 100 nM C5a for 60 minutes or 1 µM PAF for 5 minutes prior to phagocytosis of 15 µg/mL of *S. aureus* Bioparticles for 15 minutes. C5a pre-treatment led to a reduction in median phagocytosis of 33.2 % relative to control. Individual data points of each donor are shown with medians. Friedman p-value 0.042, *p < 0.05 by Dunn’s multiple comparisons test.

**B-E**: Median, IQR and outliers of the measured peptide intensity for the 2712 quantified phosphoproteins are shown for each experimental condition on each plex. Intensities are shown before (A, B) and after (C, D) median scaling normalisation of data.

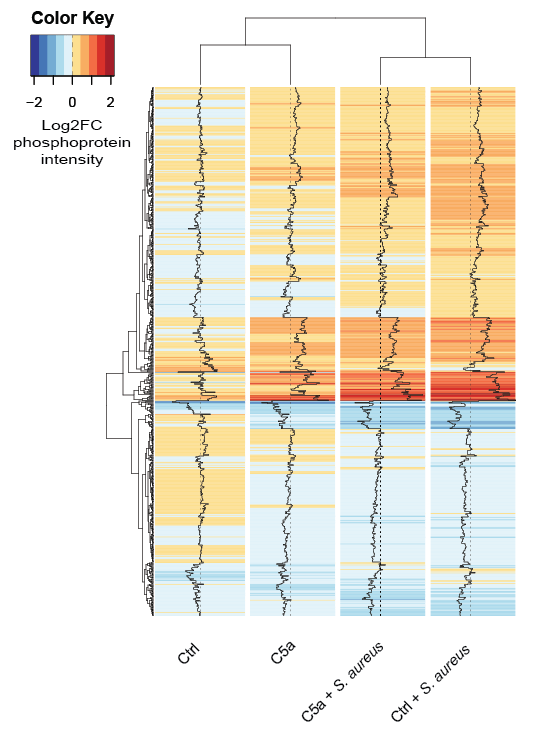

Supplemental Figure 4: *S. aureus* and C5a induce widespread changes in the neutrophil phosphoproteome, 75^th^ centile

Heatmap of phosphoprotein intensity relative to baseline (log2FC) across the 4 experimental conditions shows phosphoproteins with variance across conditions in the top 75^th^ centile with dendrograms clustered by Euclidean distance. Increased phosphoprotein expression is indicated in red, decreased in blue. Only phosphoproteins detected in all 4 donor samples were included. Protein names omitted for legibility.

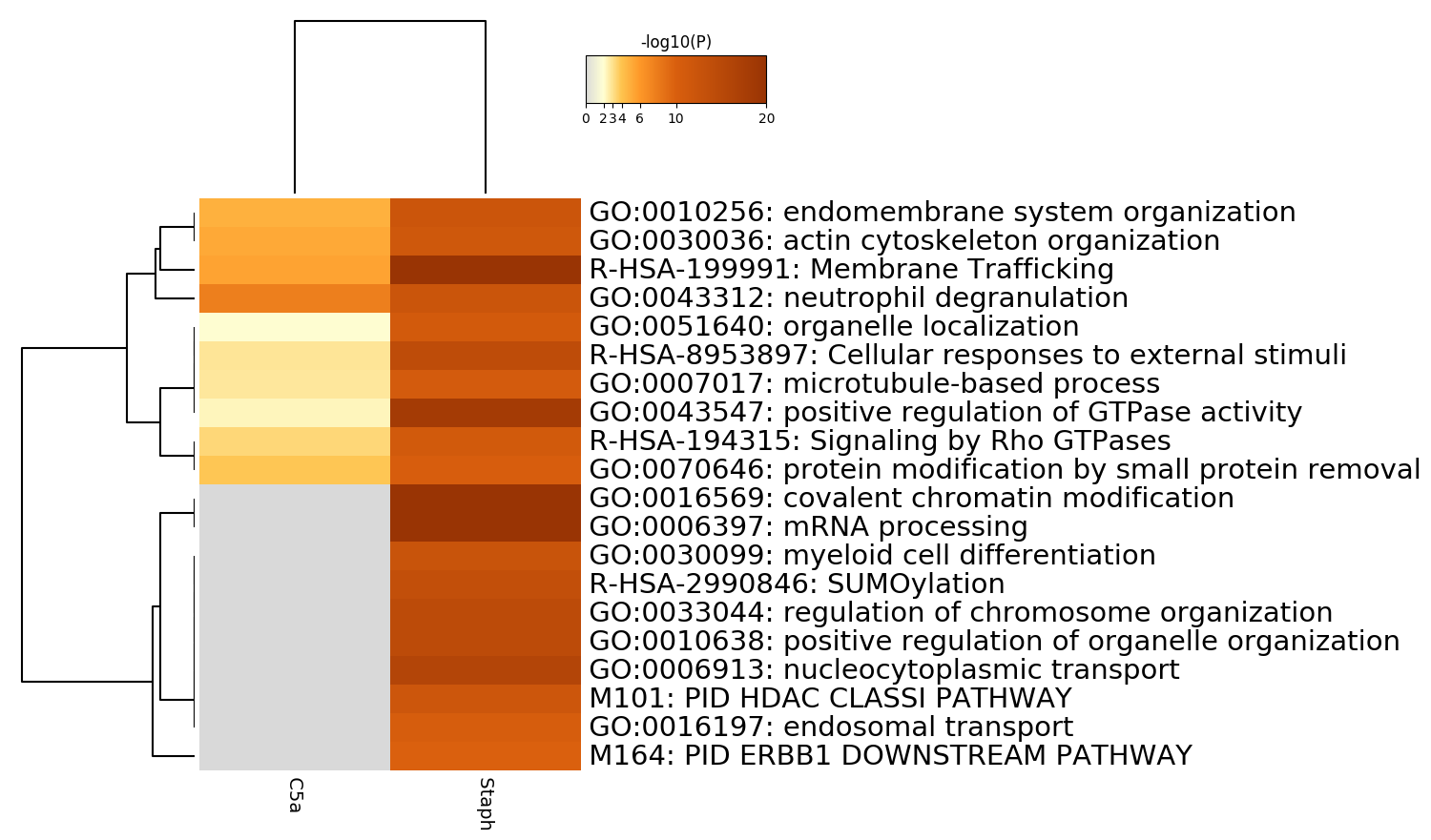
A

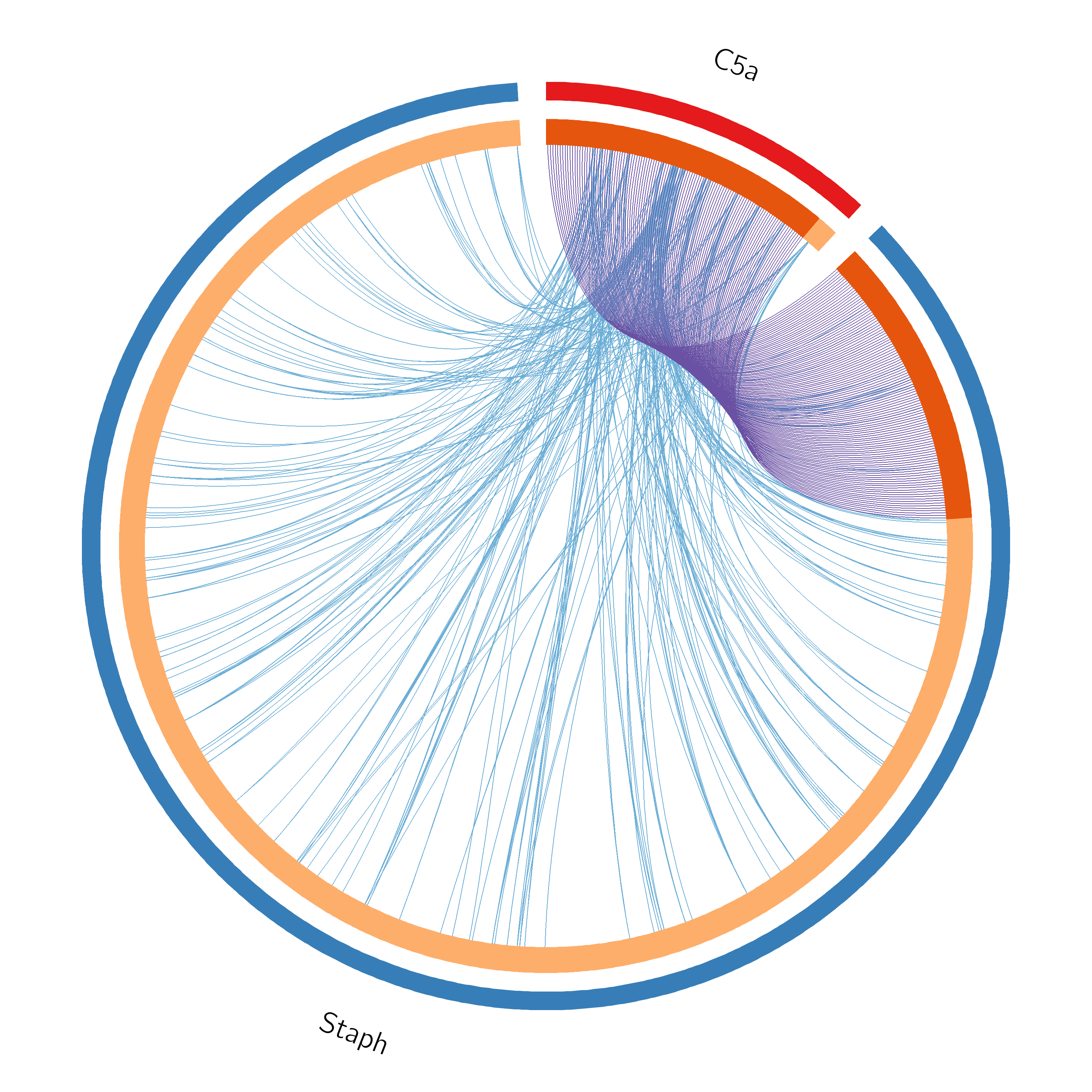

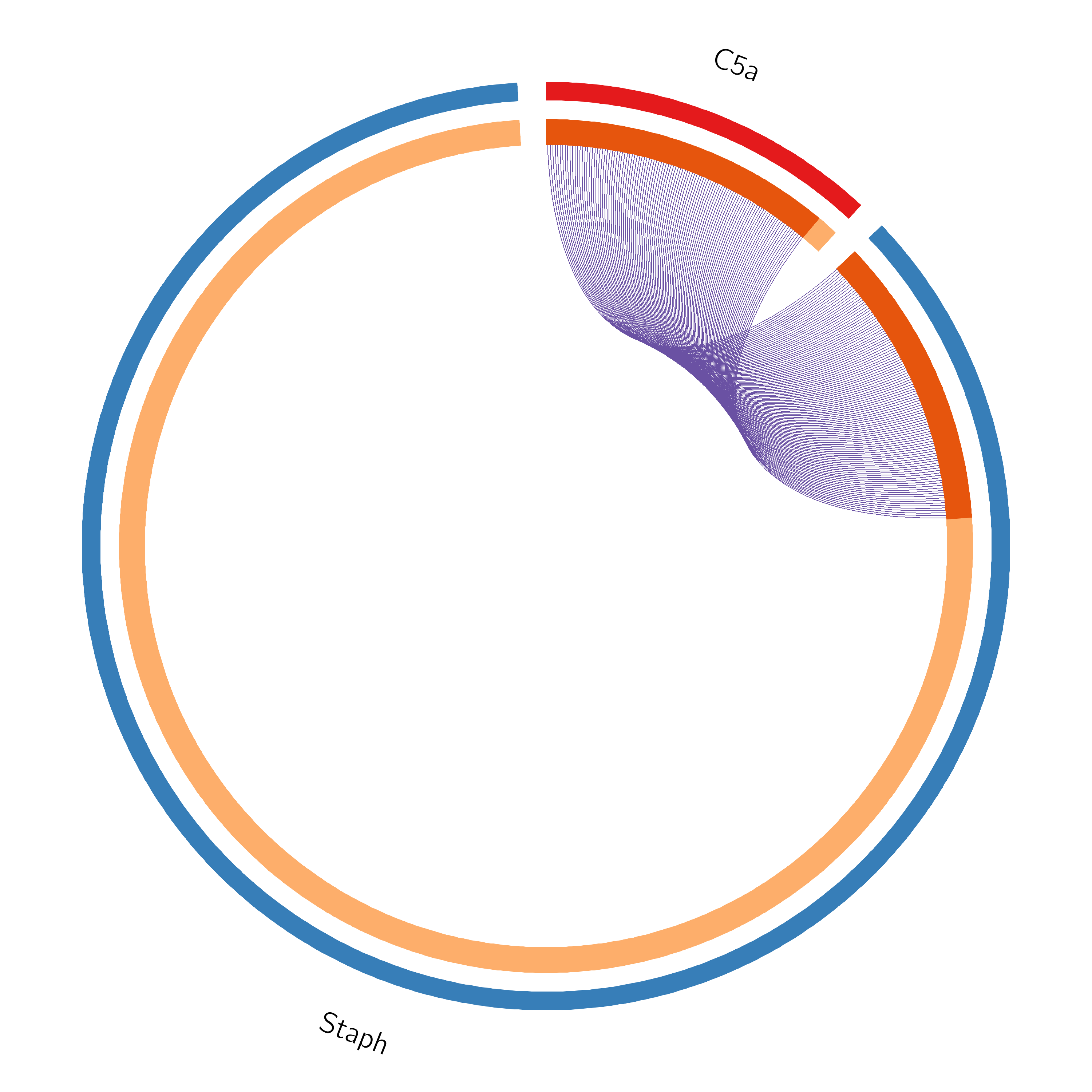
B

Supplemental Figure 5: C5a induced changes in phosphoproteins which show overlap with changes induced by *Staph aureus.*

**A:** Pathway mapping by Metascape (Zhou *et al*, 2019) demonstrates C5a-induced changes in phosphoproteins enrich pathways which are also enriched by phosphoprotein changes after *Staph. aureus* exposure.

**B**: Overlap diagram for individual phosphoproteins showing significant change under C5a and *Staph aureus* conditions, where purple lines link proteins identified in both data sets (left hand panel) and including proteins linked at the shared term level where blue lines link those proteins which belong to the same enriched ontology term (right hand panel).

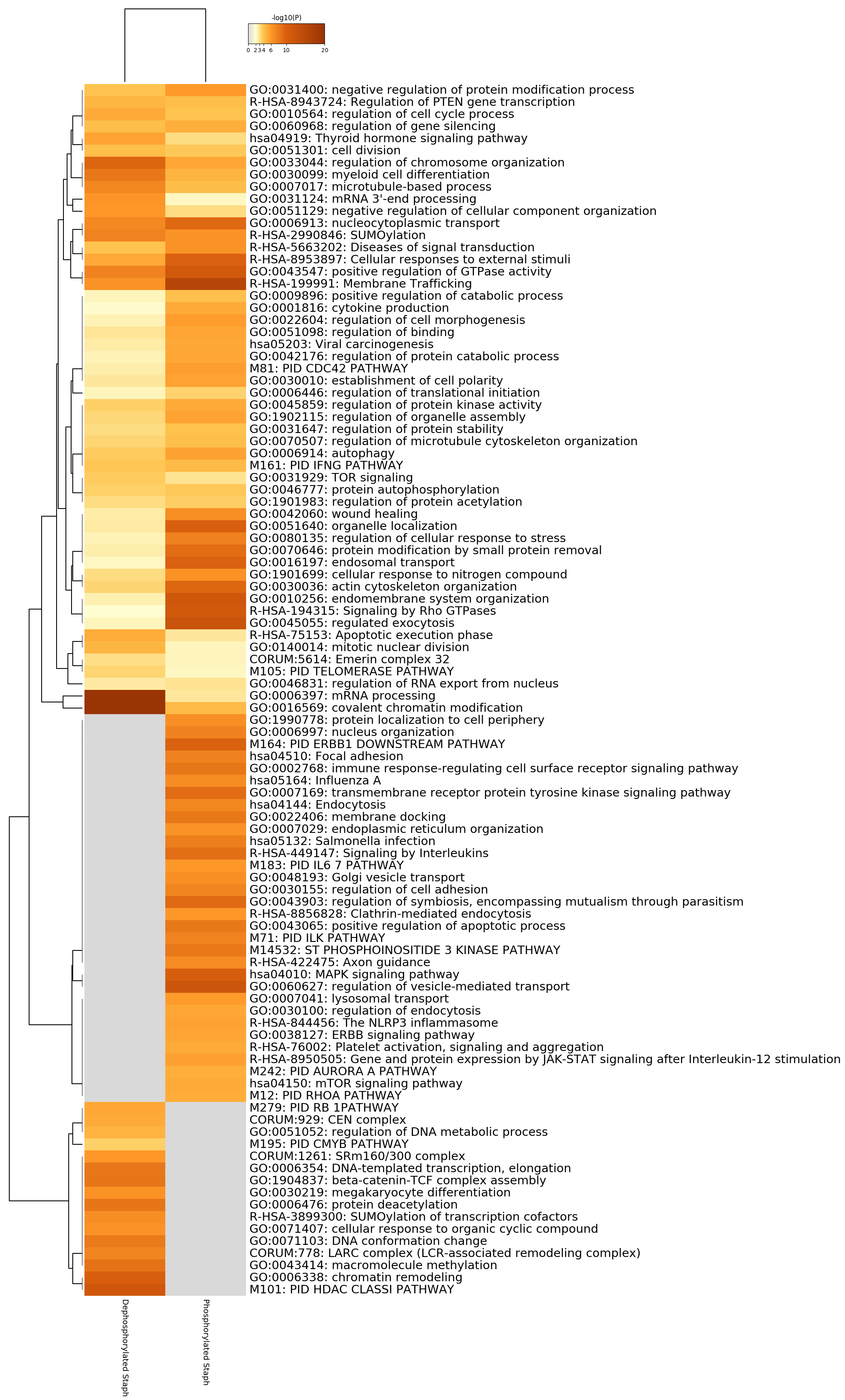

Supplemental Figure 6: Signalling pathway enriched by *S. aureus* exposure.

Metascape (Zhou *et al*, 2019) enrichment heatmap showing the top 100 functional clusters of phosphoproteins affected by *S. aureus* exposure.

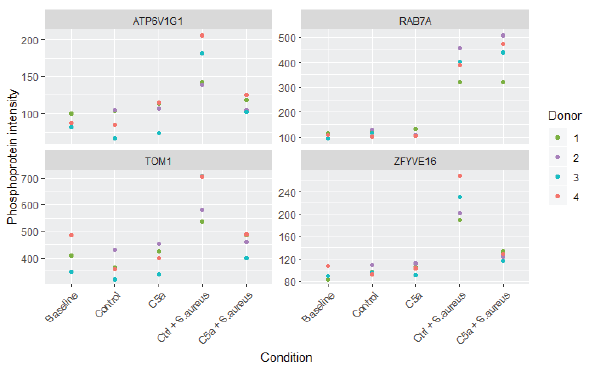

Supplemental figure 7: *S. aureus* and C5a affect signalling of key antibacterial proteins

Phosphoprotein intensities across all conditions for 4 key proteins (discussed in detail in text) are shown for each individual donor. TOM1, ZFYVE16, ATP6V1G1 and RAB7a are all involved in phagosomal maturation.

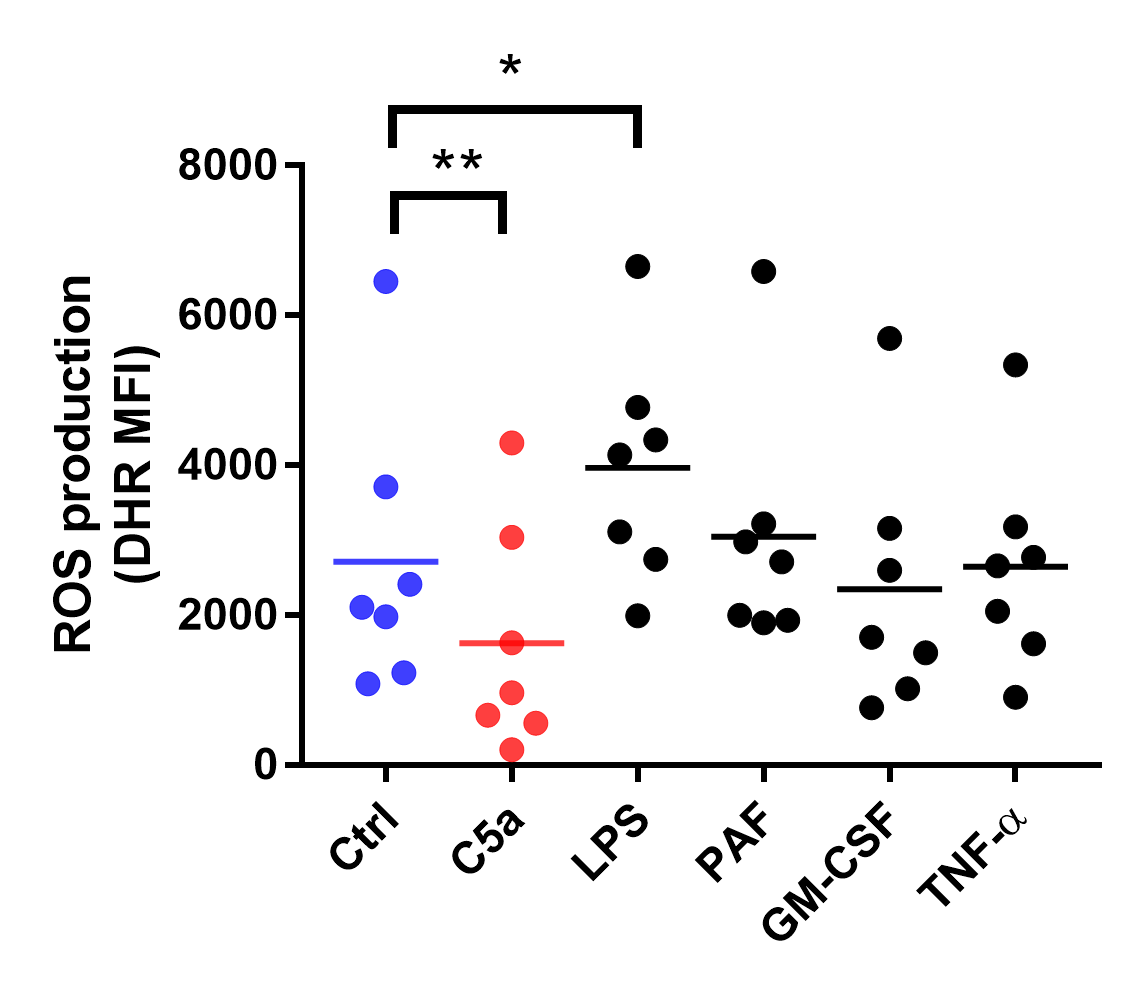

Supplemental figure 8: C5a pre-treatment reduces ROS production associated with *S. aureus* phagocytosis

Whole blood was exposed to a combined pHrodo and DHR-labelled probe for 30 minutes after pre-treatment with various agonists. Concentrations and pre-treatment durations were as follows: Ctrl (PBS) 60 minutes; C5a 300 nM, 60 minutes; LPS 100 ng/mL, 60 minutes; PAF 1 μM, 5 minutes; GM-CSF 10 ng/mL, 30 minutes; TNF-α 20 ng/mL, 30 minutes. ANOVA p-value < 0.0001 for both D and E, *p < 0.05, **p < 0.01 for Dunnett’s multiple comparisons. Data are shown as individual data points with means for n = 7 individual donors.

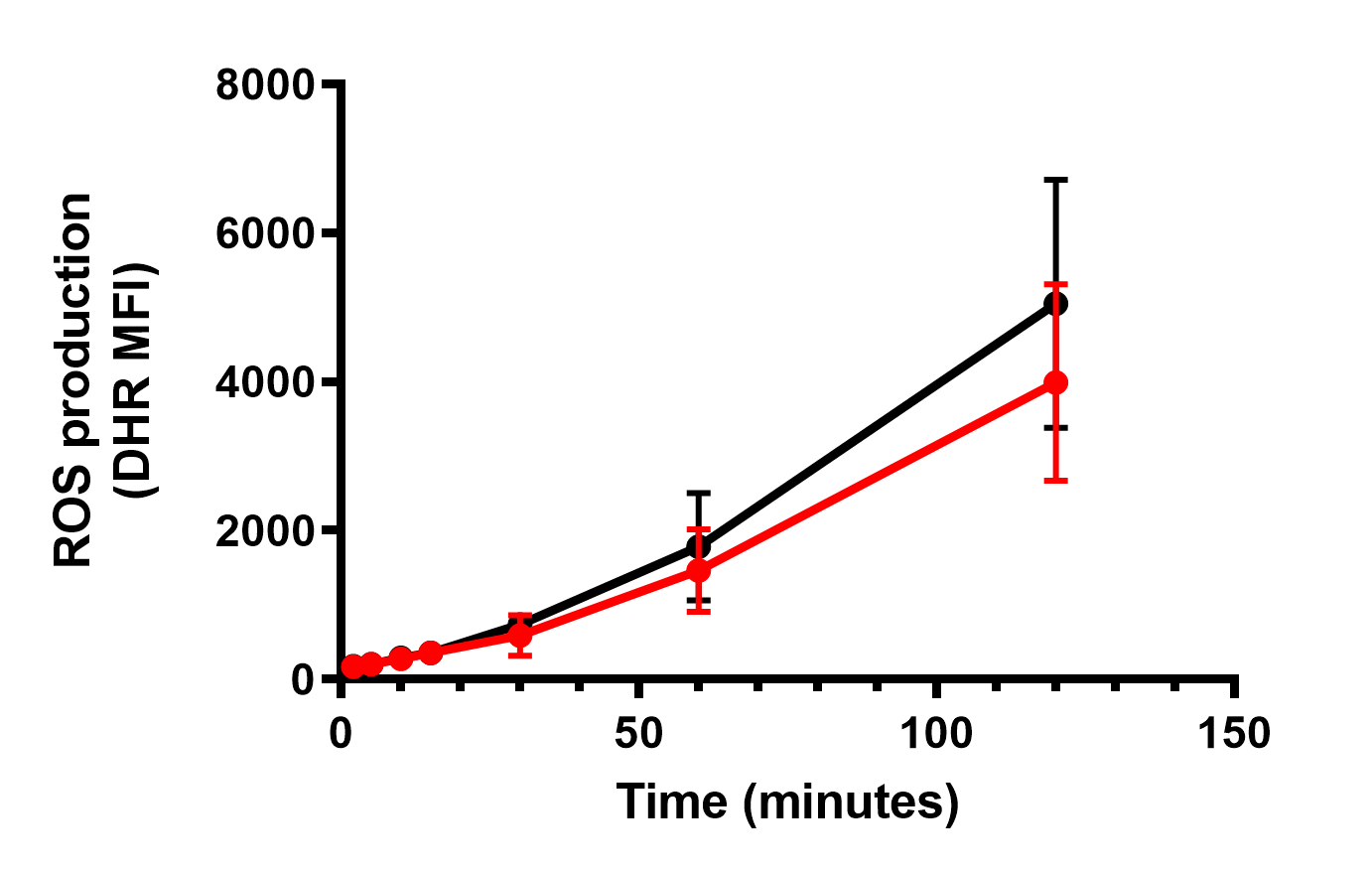

Supplemental figure 9: VPS34-IN1 did not significantly reduce phagosomal ROS production.

Whole blood was pre-treated with 1 µM VPS34-IN1 nM for 60 minutes prior to addition of the phagosomal maturation probe. Cells were allowed to phagocytose for the time points indicated. ROS production was quantified by flow cytometry. Data are shown as mean and SD of n = 5 independent experiments. No significant difference by repeated measures two-way ANOVA with Bonferroni’s multiple comparisons test.

### Supplemental methods

#### Patients and healthy donors.

Ethical permission for taking peripheral blood from healthy volunteers was obtained from the Cambridge Local Research Ethics Committee (REC reference 06/Q0108/281) with donors drawn from the healthy donor pool at Addenbrooke’s Hospital and written, informed consent obtained. Patient blood was taken under the approval granted by the North East-Newcastle & North Tyneside 2 Research Ethics Committee (REC reference: 18/NE/0036). Patients were included if they were receiving support of one or more organ systems in ICU, expected to require such support for a further 24 hours and not thought likely to die within the next 24 hours. Exclusions were pregnancy, HIV infection, haematological malignancy or immunosuppressive medication. Assent was provided a personal or nominated consultee.

#### Reagents

Reagents included purified human C5a (ComplementTech, Tyler, TX, USA), Argatroban (R&D Systems, Abingdon UK), recombinant human PAF (Tocris Bioscience, Abingdon, UK), recombinant human TNFa (R&D Systems, Abingdon, UK), recombinant human GM-CSF and *E. coli*-derived LPS (ThermoFisher Scientific, Waltham, MA, USA) and VPS34IN1 (Selleckchem, Houston, TX, USA). pHrodo™ and AF488-conjugated *E. coli* and *S. aureus* bioparticles were from ThermoFisher Scientific (Waltham, MA, USA) and DHR was from CalBiochem (Watford, UK). Antibodies used were: FITC-conjugated mouse anti-human CD88/C5aR1, clone S5/1 (BioRad, Hertfordshire, UK) and Pacific Blue-mouse anti-human CD16, clone 3G8 (Biolegend, London, UK).

#### No-wash, no-lyse whole blood assay of neutrophil phagocytosis and ROS production

This assay was adapted from a protocol provided by Life Technologies designed for use with the Attune Nxt™ acoustic focussing cytometer, see link: <https://www.thermofisher.com/content/dam/LifeTech/global/life-sciences/cellanalysis/Files/0415/Attune-No-WashLyseDetectionLeukocytes-App-Note.pdf> The initial choice of anticoagulant was informed by work from Mollnes and colleagues (Mollnes et al, 2002) which showed that direct thrombin inhibitors allowed complement activation and minimally affected leukocyte function. Blood was treated with inhibitors or priming agents as indicated in respective figure legends. After treatment, 50 µL of blood was aliquoted into 96 well plates in triplicate and a combined *S. aureus* pHrodo™/DHR probe was added at final concentrations of 15 µg/mL and 3 µM respectively. Volume was made up to 100 uL with RPMI 1640 and plates were incubated at 37 °C and 5 % CO_2_ for 30 minutes to allow phagocytosis to occur. 5 uL from each well was transferred into ice-cold RPMI 1640 media in flow cytometry tubes and stained with anti-CD16 antibody at 4C for 30 minutes. Volume was made up to 4 mL with ice-cold PBS before the cells were analysed on an Attune Nxt™ Acoustic Focusing Cytometer (LifeTechnologies, Paisely, UK). Phagocytosis was quantified by the percentage of positive neutrophils (with a sample incubated at 4°C serving as a negative control), and median fluorescent intensity of the positive cells, which were multiplied together to give a phagocytic index. In the dual target exposure two pHrodo *S. aureus* Bioparticles with different fluorescent markers were used; pHrodo green and pHrodo red were used. 50 μL of blood was aliquoted into 96 well plates in triplicate and the first Bioparticle *S. aureus* pHrodo red was added at a final concentration indicated in the figure legend and incubated for 30 minutes at 37 °C and 5 % CO2 to allow phagocytosis to occur. The second Bioparticle *S. aureus* pHrodo green was then added and cells were allowed to phagocytose as before. Samples were then stained for CD16 and analysed as above.

#### Bacterial killing assay – whole blood

Methicillin-sensitive *Staphylococcus aureus* (MSSA) bacteria (strain ASASM6, kind gift from Prof Gordon Dougan, University of Cambridge) were sub-cultured in 10 mLs of Luria broth (LB) overnight, then grown to early log-phase. Human blood was collected into argatroban (300 µg/mL) and aliquoted into Eppendorf tubes (50 µL). 10 µL of bacterial suspension was added and blood was incubated on a shaker at 200 rpm, at 37˚C for 1 hour. Human cells were lysed by addition of 450 µL pH 11 distilled water for 3 minutes. Samples were vortexed for 15 seconds and lysates diluted 1:1000 with LB then 100 µL of diluted sample was plated on blood agar plates. Plates were incubated overnight at 37˚C and colonies were counted by visual inspection the next morning.

#### Bacterial killing assay – purified neutrophils

Methicillin-sensitive *Staphylococcus aureus* (MSSA) bacteria (strain ASASM6) were sub-cultured in 10 mLs of Luria broth (LB) overnight, then grown to early log-phase. In the interim, isolated neutrophils were prepared at a concentration of 5 million/mL in IMDM before pre-incubation with 100nM C5a or vehicle control for 60 minutes before exposure to MSSA at an. average multiplicity of infection (MOI) of 10:1, for one hour at 200 rpm, 37 ^o^C. Samples were then treated with distilled water for 3 minutes to lyse the neutrophils without adversely affecting MSSA viability and serial dilutions plated on blood agar for overnight culture and visual counting of colonies.

#### Proteomics

#### Cell lysis and trypsin digestion

Cells were lysed in freshly prepared ice-cold lysis buffer (0.5 % SDS, 0.1 M TEAB containing 1 X HALT protease and phosphatase inhibitors). Cell suspensions were incubated at 90 °C for 5 minutes and sonicated twice for 20 seconds (EpiShear™ Probe Sonicator, Active Motif). The insoluble fraction was removed by centrifugation at 20 000 g for 10 minutes at 4 °C. Lysate protein concentration was determined by Bradford Protein Assay (Bio-Rad, Quick Start).

100 µg of protein per sample was reduced in 5 mM Tris(2-carboxyethyl)phosphine‎ (TCEP) for one hour at 60 °C, which was followed by alkylation of cysteines with 10 mM methyl methanethiosulfonate in the dark at room temperature. Lysates were then diluted 1:10 with 0.1 M TEAB before trypsin was added at a 1:30 trypsin:protein ratio by mass. Digestion was carried out overnight at room temperature.

#### Tandem mass tag labelling

Tandem mass tag (TMT) reagents (0.8 mg) were reconstituted in 40 µl anhydrous acetonitrile and peptide samples were labelled according to the manufacturer’s protocol. Following incubation at room temperature for one hour, the reaction was quenched with hydroxylamine to a final concentration of 5% (v/v) for 15 min at room temperature. The TMT-labelled samples were pooled at 1: 1: 1: 1: 1: 1: 1: 1: 1: 1: 1 across the 11 samples. The pooled sample was vacuum centrifuged to dryness and subjected to basic pH reversed-phase (bRP) fractionation.

#### Off-line reverse phase fractionation at basic pH

The TMT mixture was fractionated on a Dionex Ultimate 3000 system (Thermo Fisher Scientific) at high pH. The labelled peptide mixture was reconstituted in 20 mM ammonium hydroxide in water (pH 10) and subjected to a 45 minute linear gradient from 4.5% to 45% acetonitrile in 20 mM ammonium hydroxide (pH 10) at a flow rate of 0.2 ml/min over an XBridge C18 column Reversed-Phase (3.5 µm particles, 2.1 mm ID, 150 mm in length; Waters). The peptide mixture was fractionated into a total of 48 fractions. 15% (v/v) of each fraction was separated into a new tube and was used for full proteome analysis. The remaining volume in each fraction will be subjected to phosphopeptide enrichment. All fractions were dried on a centrifugal vacuum concentrator.

#### Phosphopeptide enrichment

The initial 48 fractions were consolidated into 14 and submitted to phosphopeptide enrichment by using the High-Select™ Fe-NTA Phosphopeptide Enrichment Kit according to the manufacturer’s protocol. Eluates containing the phosphopeptides were dried via vacuum centrifugation and reconstituted in 10 µl of 0.1 % formic acid for liquid chromatography and tandem mass spectrometry (LC-MS/MS) processing.

#### LC-MS/MS analysis of full proteome fractions

The initial 48 fractions were pooled into a final number of 32 fractions. Each fraction was reconstituted in 10 µl of 0.1% formic acid and 5 µl were analysed on a Dionex Ultimate 3000 UHPLC system coupled with an Orbitrap Fusion Lumos mass spectrometer (Thermo Fisher Scientific, San Jose, CA). Samples were loaded on an Acclaim PepMap 100, 100 μm × 2 cm C18, 5 μm, 100 Å trapping column with the ulPickUp injection method at a loading flow rate of 5 μL/min for 10 min. For peptide separation, an EASY-Spray analytical column 75 μm × 25 cm, C18, 2 μm, 100 Å column was used for multi-step gradient elution at a flow rate of 300 nL/min. Mobile phase (A) was composed of 2 % acetonitrile, 0.1 % formic acid; mobile phase (B) was composed of 80 % acetonitrile, 0.1 % formic acid. Peptides were eluted using a gradient as follows: 0 - 10 min, 5 % mobile phase B; 10 – 95 min, 5 – 45% mobile phase B; 95 -100 min, 45% - 95% B; 100 - 108 min, 95% B; 108 – 110min, 95% - 5% B; 110 – 120 min, 5% B.

Data-dependent acquisition began with an MS survey scan in the Orbitrap (380 – 1500 m/z, resolution 120,000 full width half maximum (FWHM), automatic gain control (AGC) target 3E5, maximum injection time 100 ms). The top ten precursors were then selected for MS2/MS3 analysis. MS2 analysis consisted of: collision-induced dissociation (CID), quadrupole ion trap analysis, AGC target 1E4, normalized collision energy (NCE) 35, q-value 0.25, maximum injection time 35 ms, an isolation window at 0.7, and a dynamic exclusion duration of 45 seconds. Following acquisition of MS2 spectrum, MS3 precursors were fragmented by high energy collision-induced dissociation (HCD) using 10 frequency notches and analysed in the Orbitrap (resolution 50,000 FWHM, AGC target 5E4, NCE 55, maximum injection time 86 ms, and isolation window at 0.7.

#### LC-MS/MS analysis of phosphopeptide-enriched fractions

All the fractions (10 µl) were analysed on a Dionex Ultimate 3000 UHPLC system coupled to a Q-Exactive HF mass spectrometer (Thermo Fisher Scientific, San Jose, CA). Samples were loaded on an Acclaim PepMap 100, 100 μm × 2 cm C18, 5 μm, 100 Å trapping column with the ulPickUp injection method at a loading flow rate of 5 μL/min for 10 min. For peptide separation, an EASY-Spray analytical column 75 μm × 25 cm, C18, 2 μm, 100 Å column was used for multi-step gradient elution at a flow rate of 300 nL/min. Mobile phase (A) was composed of 2 % acetonitrile, 0.1 % formic acid, 5 % dimethyl sulfoxide (DMSO); mobile phase (B) was composed of 80 % acetonitrile, 0.1 % formic acid, 5 % DMSO. Peptides were eluted using a gradient as follows: 0 - 10 min, 5 % mobile phase B; 10 – 95 min, 5 – 45 % mobile phase B; 95 -100 min, 45 % - 95% B; 100 - 108 min, 95 % B; 108 – 110min, 95 % - 5 % B; 110 – 120 min, 5 % B.

Data-dependent acquisition began with an MS survey scan in the Orbitrap (400 – 1600 m/z, resolution 60,000 FWHM, AGC target 3E6, maximum injection time 100 ms). The top 10 precursors were then isolated and fragmented with CID. MS2 analysis consisted of: resolution 30,000 FWHM, AGC target 2E4, NCE 33, maximum injection time.

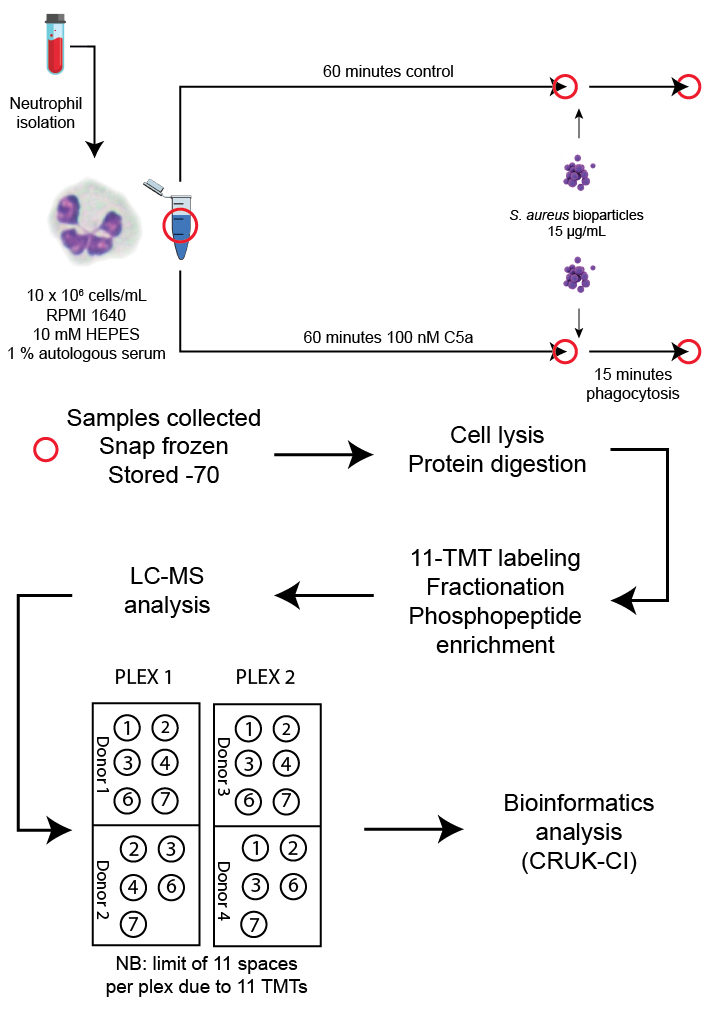
 100 ms, MS2 isolation window 2.0 m/z.

Supplemental figure 10: Schematic of phosphoproteomic workflow

Neutrophils were purified as described and suspended in RPMI 1640 media supplemented with 10 mM HEPES and 1% AS. Red circles indicate instances where aliquots of cells were removed for analysis. An untreated aliquot was removed and snap frozen and cells were then treated with 100 nM C5a, or control for 60 minutes. Post-treatment, pre-phagocytosis aliquots were removed and snap frozen, and pHrodo™ *S. aureus* Bioparticles were added; cells were allowed to phagocytose for 15 minutes before the final aliquots were removed and snap frozen. Phagocytosis was measured at the end of the procedure by flow cytometry of remaining (non-frozen) sample. Samples were batched until all 22 conditions were ready for analysis. Samples were transported to CRUK-CI on dry ice, lysed and digested as specified. The phosphoproteomic workflow included TMT labelling, fractionation and enrichment of phosphopeptides prior to LC-MS analysis in two separate batches or plexes with the 22 samples from four donors distributed across them as shown.
